# Quantifying dispersal variability among nearshore marine populations

**DOI:** 10.1101/2020.09.17.299941

**Authors:** Katrina A. Catalano, Allison G. Dedrick, Michelle R. Stuart, Jonathan B. Puritz, Humberto R. Montes, Malin L. Pinsky

## Abstract

Dispersal drives diverse processes from population persistence to community dynamics. However, the amount of temporal variation in dispersal and its consequences for metapopulation dynamics is largely unknown for organisms with environmentally driven dispersal (e.g., many marine larvae, arthropods, and plant seeds). Here, we quantify variation in the dispersal kernel across seven years and monsoon seasons for a common coral reef fish, *Amphiprion clarkii*, using genetic parentage assignments. Connectivity patterns varied strongly among years and seasons in the scale and shape but not in the direction of dispersal. This interannual variation in dispersal kernels introduced temporal covariance among dispersal routes with overall positive correlations in connections across the metapopulation that may reduce stochastic metapopulation growth rates. The extent of variation in mean dispersal distance observed here among years is comparable in magnitude to the differences across reef fish species. Considering dispersal variability will be an important avenue for further metapopulation and metacommunity research across diverse taxa.

## Introduction

Metapopulation structure is common across diverse taxa, from arthropods and fungal spores to plant seeds and marine larvae (Lett, Barrier, & Bahlali, 2020). In a metapopulation, connectivity through dispersal drives a wide range of ecological and evolutionary processes (R. K. Cowen, Paris, & Srinivasan, 2006; Gotelli, 1991; Hanski, 1994), including population persistence (Hastings & Botsford, 2006; Holyoak & Lawler, 1996), genetic population differentiation and local adaptation (Kawecki & Ebert, 2004; Taylor & Hellberg, 2003), and community assembly and dynamics (Gaines & Roughgarden, 1985; Leibold et al., 2004). For conservation and management, connectivity is an important consideration in the design of protected areas and for invasive species management (Kough, Paris, & Butler IV, 2013; Saura, Bodin, & Fortin, 2014; Tamburello, Ma, & Côté, 2019; Ying, Chen, Lin, Gao, & Quinn, 2011).

Dispersal, however, may vary strongly through time, and this variation is predicted to have important consequences for metapopulation dynamics (Snell et al., 2019; Snyder, Paris, & Vaz, 2014; James R. Watson, Kendall, Siegel, & Mitarai, 2012; Williams & Hastings, 2013), competition and predation (Wright, Muller-Landau, Calderón, & Hernandéz, 2005), invasion dynamics (Ellner & Schreiber, 2012), community assembly (Matias, Mouquet, & Chase, 2013), and responses to climate change (Pearson & Dawson, 2005), among other processes. For a metapopulation to persist, the metapopulation growth rate must be high enough for individuals to replace themselves in a lifetime (Hastings & Botsford, 2006). Theory predicts that when connectivity between patches fluctuates and is positively correlated between pairs of patches, the long-term growth rate of the metapopulation will be lower than the growth rate calculated using time-averaged connectivity (Bani, Fortin, Daigle, & Guichard, 2019; Snyder et al., 2014; James R. Watson et al., 2012). Positively correlated connectivity can occur when individuals from multiple subpopulations are entrained in the same flow regime (e.g., wind or water). In contrast, if dispersal from different patches is anti-correlated by reversals in flow or other mechanisms, metapopulation growth rates may instead be enhanced (Williams & Hastings, 2013). To date, these questions have been addressed primarily with dispersal simulations (Bani et al., 2019; Castorani et al., 2017; Heydel, Cunze, Bernhardt-Römermann, & Tackenberg, 2014; James R. Watson et al., 2012), but empirical observations are necessary to understand variability in both the distance and direction of dispersal through time. Though there has been a surge in research quantifying dispersal (Abesamis et al., 2017; D’Aloia, Bogdanowicz, Majoris, Harrison, & Buston, 2013; Planes, Jones, & Thorrold, 2009; Salles et al., 2015), empirical measurements of temporal variation in dispersal are a critical step toward understanding how connectivity affects the ecological and evolutionary dynamics of metapopulations.

Variable dispersal is likely to be particularly important in marine metapopulations because the pelagic larvae of many marine organisms are transported by complex and often chaotic ocean currents that can drive stochastic connectivity (Mitarai, Siegel, & Winters, 2008; Siegel et al., 2008). Transient oceanographic features such as mesoscale eddies, surface fronts, and alongshore jets vary over daily, seasonal, and annual time scales and shape dispersal patterns. Simulations of dispersal demonstrate that this variability in flow can generate both spatially and temporally heterogeneous dispersal (R. K. Cowen et al., 2006; Mitarai et al., 2008; Siegel et al., 2008; J. R. Watson et al., 2010). Indeed, spatial genetic structure, which is influenced by dispersal, is known to vary temporally (Toonen & Grosberg, 2011). As a result, marine larval dispersal is expected to vary temporally, although the empirical character and extent of this variation is unclear.

Only a few studies have tackled the logistically difficult task of directly measuring larval dispersal in more than one time period. These contributions have demonstrated empirically that dispersal can vary seasonally within a year (Abesamis et al., 2017; Carson, López-Duarte, Rasmussen, Wang, & Levin, 2010) and among years (Almany et al., 2017; Andrew & Ustin, 2010; Berumen et al., 2012; Hogan, Thiessen, Sale, & Heath, 2012; Ran Nathan, Safriel, Noy-Meir, & Schiller, 2000), but have been limited to two to three years of sampling and have not yet been contextualized to understand the implications for metapopulation persistence. Connectivity is often inferred from the dispersal distances between the origin and recruitment locations of individuals, often through genetic parentage analysis (Berumen et al., 2012; D’Aloia et al., 2013; Geoffrey P Jones, Milicich, Emslie, & Lunow, 1999; Swearer, Caselle, Lea, & Warner, 1999). With these data, an isotropic probability density function, called a dispersal kernel, can be statistically fit to data to describe how the strength of dispersal varies with distance (Bode, Williamson, Harrison, Outram, & Jones, 2017). The dispersal kernel then characterizes the frequency of dispersal and subsequent recruitment across spatial scales. To understand how variation in dispersal impacts metapopulation dynamics, there is a need to quantify a comprehensive time series of dispersal kernels across years and measure this variation in dispersal through time.

Our objectives here were to infer annual and seasonal dispersal using genetic parentage assignment across seven years (2012-2018) in a population of yellowtail anemonefish, *Amphiprion clarkii*. We quantified interannual and seasonal variation in the dispersal kernel to understand changes in the dispersal distances. To consider the implications for metapopulation persistence, we also tested for changes in the direction of dispersal and measured correlations in connectivity routes through time.

## Material and Methods

### Study system

We sampled yellowtail anemonefish (*Amphiprion clarkii*) on 19 reef patches (sites) along approximately 30 km of coastline in Ormoc Bay, Leyte, Philippines (Fig. 1). The Philippines has a tropical climate with seasonal monsoon cycles. The Northeast Monsoon occurs November-May and is characterized by low rainfall. The Southwest Monsoon brings heavier rainfall June-October, also coinciding with the tropical cyclone season July-October (Lau & Yang, 1997). *A. clarkii* occupy sea anemones, which makes it relatively easy to find individuals in the field. Because *A. clarkii* remain in their home anemone after settlement, the distance between a parent and a recruit is the dispersal distance. They are protandrous hermaphrodites, and the largest two adults in an anemone breed (Haruki Ochi, 1989). Benthic eggs develop for six days before hatching into the water column (Holtswarth, Jose, Montes, Morley, & Pinsky, 2017). Dispersal occurs during the pelagic larval stage for 8-10 days before settlement to an anemone (Thresher, Colin, & Bell, 1989). *A. clarkii* spawn on a lunar cycle all year long, but produce the most eggs from November to May (Holtswarth et al., 2017). Generation time is likely to be around five years based on growth and maturity in this species (Moyer, 1986; H. Ochi, 1986).

**Figure 1.**
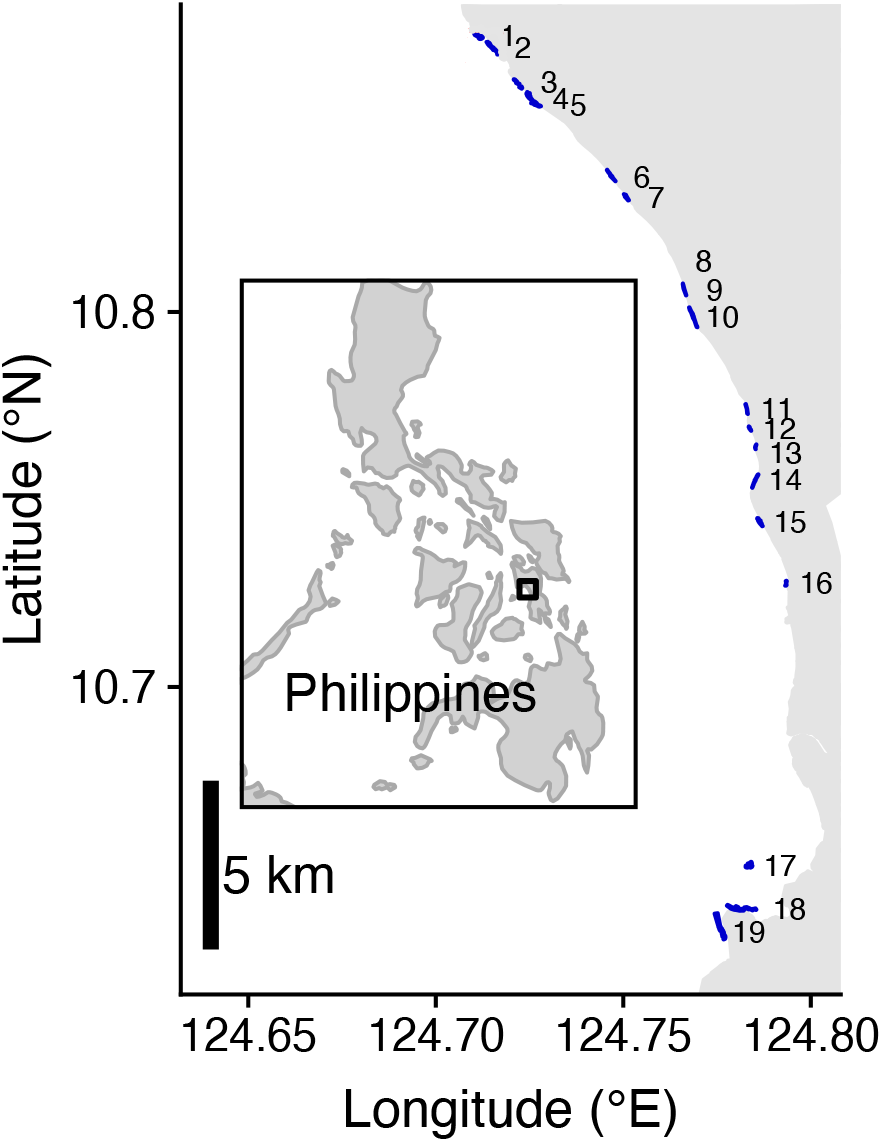
Map of the study region in Ormoc Bay, Leyte, Philippines. Study region covers about 30 km of coastline, including 19 habitat patches: Palanas (1), Wangag (2), North Magbangon (3), South Magbangon (4), Cabatoan (5), Caridad Cemetery (6), Caridad Proper (7), Hicgop South (8), Sitio Tugas (9), Elementary School (10), Sitio Lonas (11), San Agustin (12), Poroc San Flower (13), Poroc Rose (14), Visca (15), Gabas (16), Tomakin Dako (17), Haina (18), Sitio Baybayon (19).

### Genetic parentage analysis

Tissue samples were collected annually from 2012-2018 (see Supporting Information). We prepared genomic libraries for sequencing with a ddRADseq protocol (Peterson, Weber, Kay, Fisher, & Hoekstra, 2012) and genotyped 791 recruits and 1,729 adult *A. clarkii* at 1,340 Single Nucleotide Polymorphisms (SNPs) using the bioinformatics pipeline dDocent (Puritz, Hollenbeck, & Gold, 2014) (see Supporting Information). We then used Colony2 to identify parent-offspring matches (Wang, 2012). Colony2 uses maximum likelihood to jointly estimate parentage and sibship relationships from SNP data. We ran Colony2 for each year by including only recruits sampled in that year. We cumulatively pooled adults across years so that once an adult was sampled, it was included as a possible parent in the analysis of that and all subsequent years (Table S1). We retained parentage assignments found in the maximum likelihood configuration.

### Fitting dispersal kernels

We used maximum likelihood to fit probabilistic dispersal kernels (Bode et al., 2017) to each year’s parentage results separately, to the combined results from 2012-2018, and to seasonal cohorts of recruits across 2012-2018 (see Supporting Information). We fit generalized Gaussian kernels that describe the relative probability density of dispersal (*p*) from the natal patch *i* to patch *j* with increasing distance (*d*) as:

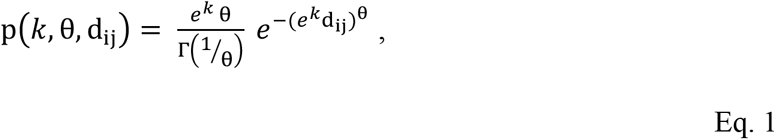

where Γ denotes the gamma function. The generalized Gaussian is defined by a scale parameter *k* that sets the dispersion of the distribution and a shape parameter θ that affects its kurtosis (how much of the density is contained in the tail). At a value of θ = 1, the kernel is Laplacian and leptokurtic, exhibiting exponential decline with increasing distance and a fat tail. In contrast, θ = 2 is a Gaussian kernel, while θ = 3 is a platykurtic Ribbens function with a broad central peak and a thin tail.

We calculated mean dispersal distance, as well as the distance at which 50% (median) and 90% of larvae are expected to recruit for each kernel. Then, we calculated a distribution of mean and median distances from 1,000 parameter sets of *k* and θ drawn with replacement from each kernel 2D likelihood surface, weighting parameter sets by the difference between the log-likelihood value and the maximum log-likelihood value (parameter sets closer to the maximum log-likelihood are thus more likely to be drawn). Finally, we integrated the kernels across the width of our study region (30 km) to estimate the proportion of recruits retained in the metapopulation.

To examine variation in kernel shape, we quantified the excess kurtosis (the density in the tails relative to a Gaussian kernel) using the following equation (Clark et al., 1998),

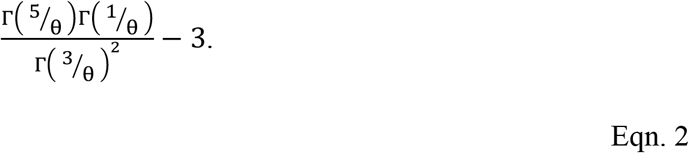

Negative values indicate a platykurtic (thin-tailed) distribution, meaning that long distance dispersal events are rare, and positive values indicate a leptokurtic (fat-tailed) distribution, meaning long-distance dispersal events are relatively common.

We also explored whether dispersal is variable on seasonal (monsoonal) timescales by fitting kernels separately to recruits likely to have dispersed during the Northeast Monsoon or the Southwest Monsoon. We generally sampled in late May-June of each year (the end of the Northeast Monsoon season, see Supporting Information), and so we did not sample directly in each season. Rather, we used an estimate of fish growth rates to partition sampled recruits into Northeast Monsoon or Southwest Monsoon cohorts. Recruits less than 3.5 cm were likely to have dispersed in the preceding five months during the Northeast Monsoon (between November and June). In contrast, recruits greater than 4.5 cm but less than 6 cm were likely to be 6 to 11 months old and to have dispersed during the Southwest Monsoon.

We tested whether the observed interannual and seasonal variation in kernel fits was greater than expected from a null model that simulated no differences among years or seasons. We randomly shuffled the years in which recruits were observed, creating a seven-year time series that broke apart any relationship between year and dispersal kernel. We fit kernels to each of the seven years, calculated the standard deviation of *k* across years and calculated the coefficient of variation for θ, mean dispersal distance, and median across years. We then repeated the reshuffling 1,000 times. We compared these null distributions of temporal variation to the empirical estimates. We used the same approach for seasonal dispersal variation.

Finally, for both the annual and seasonal kernel fits, we explored the sensitivity of our results to the proportion of the population sampled. We down-sampled the all-years data (2012-2018 for both assigned and unassigned recruits) at proportions ranging from 0.01-1.0 and fit a kernel to each of *n* = 100 samples. We then plotted the relationship between the proportion of data sampled and either *k* or θ.

### Variation in dispersal direction

Next, we tested whether average dispersal direction changed among years. We fit generalized linear models (GLMs) with binomial errors where the response variable indicated whether a parentage match was observed (1) or not (0) for each pairwise site-to-site connection (dispersal route) each year (D’Aloia et al., 2013). Predictor variables included the distance between sites, the sampling effort at the source (adult) and destination (recruit) sites, the source and destination site size (number of anemones), the sampling year, and the direction from source to destination (North = +1, same site = 0, South =−1). When larvae recruit to their natal (source) site, we refer to this as self-recruitment. We calculated sampling effort as the proportion of anemones sampled at a site in each year (see Supporting Information). To test directional switching of dispersal among years, we included an interaction term between year and directionality. We used Akaike information criterion (AIC) to sequentially remove terms from the full model. We used the same approach to explore variation in direction between seasons, using season as a predictor variable rather than year.

### Correlation among dispersal routes

Connectivity between pairs of sites (dispersal routes) underlies metapopulation dynamics (Hanski, 1994; Hastings & Botsford, 2006; Roughgarden & Iwasa, 1986). To examine temporal variation in each dispersal route, we discretized the annual kernels according to the known widths of and distances between sites, creating a connectivity matrix for each year where each cell (a single dispersal route) represented the probability of dispersal from a source site (column) to a destination site (row). We then calculated the coefficient of variation for each dispersal route across the seven years.

Finally, because the effects of connectivity variation depend on how connectivity covaries among routes (Snyder et al., 2014; James R. Watson et al., 2012; Williams & Hastings, 2013), we calculated Pearson’s correlation for each pair of routes across the seven years (Snyder et al., 2014). We used Ward’s hierarchical clustering to identify groups of routes that maximized similarity of the correlation coefficient within the group. We chose the number of clusters so that the within-cluster sum of squares showed little further decrease with a larger number of clusters.

## Results

### Genetic parentage analysis

The parentage analysis identified 71 parent-offspring matches from 791 recruits and 1,729 adults for an overall assignment rate of 9% (Table S2). We also identified parents for 11 of 132 offspring that likely dispersed during the Northeast Monsoon and for 35 of 428 offspring from the Southwest Monsoon (Table S3). Of the assigned offspring, 28 were assigned to both parents. When both parents were identified, they had always been sampled on the same anemone. Overall, the frequency of observed dispersal events generally decreased as distance increased (Fig. S1). Across all years, 27 individuals dispersed northward, 23 dispersed southward, and 21 recruited to their natal site (Fig. 2).

**Figure 2.**
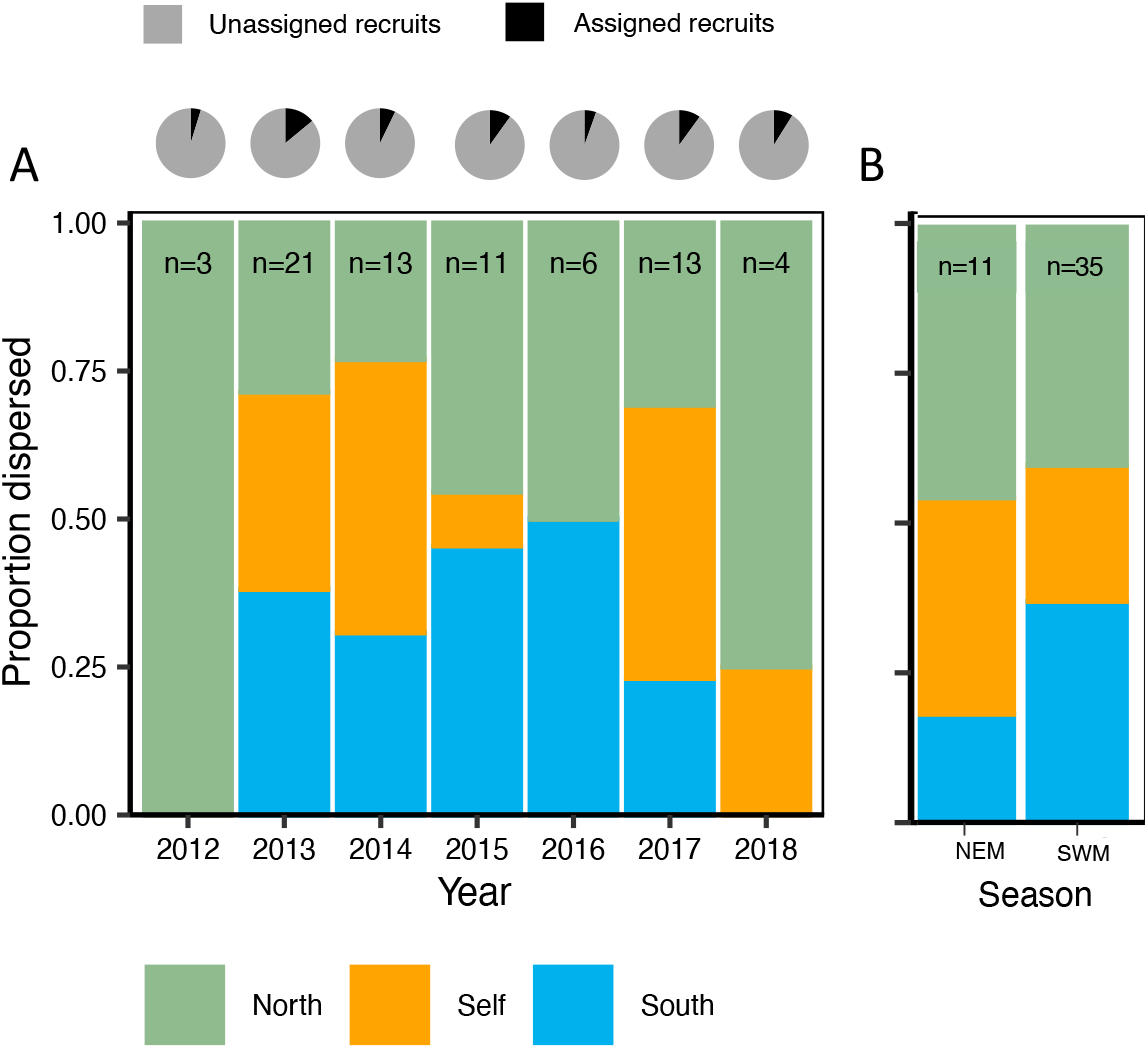
The proportion of observed dispersal events that were northwards (green), southwards (blue), or self-recruiting (orange) relative to the natal site in each year (A) and season (B). The pie charts represent the proportion of sampled recruits in each year that were assigned (or not) to sampled parents.

### Dispersal kernels

The best fit parameter sets describing the scale (*k*) and shape (θ) of the dispersal kernels differed among years (Fig. 3, Table S2, Table S4), though the bivariate likelihood surfaces overlapped for some years (Fig. S2). Dispersal probabilities dropped especially quickly with distance in 2013, 2014, 2015, and 2017, but declined more slowly in 2012, 2016, 2018, and the average kernel (Fig. 3). Annual variation in scale (SD = 2.80) was significantly greater than expected from the null model (*p* = 0.001), as was shape θ (CV = 1.33, *p* = 0.001, Fig. S3A,B).

**Figure 3.**
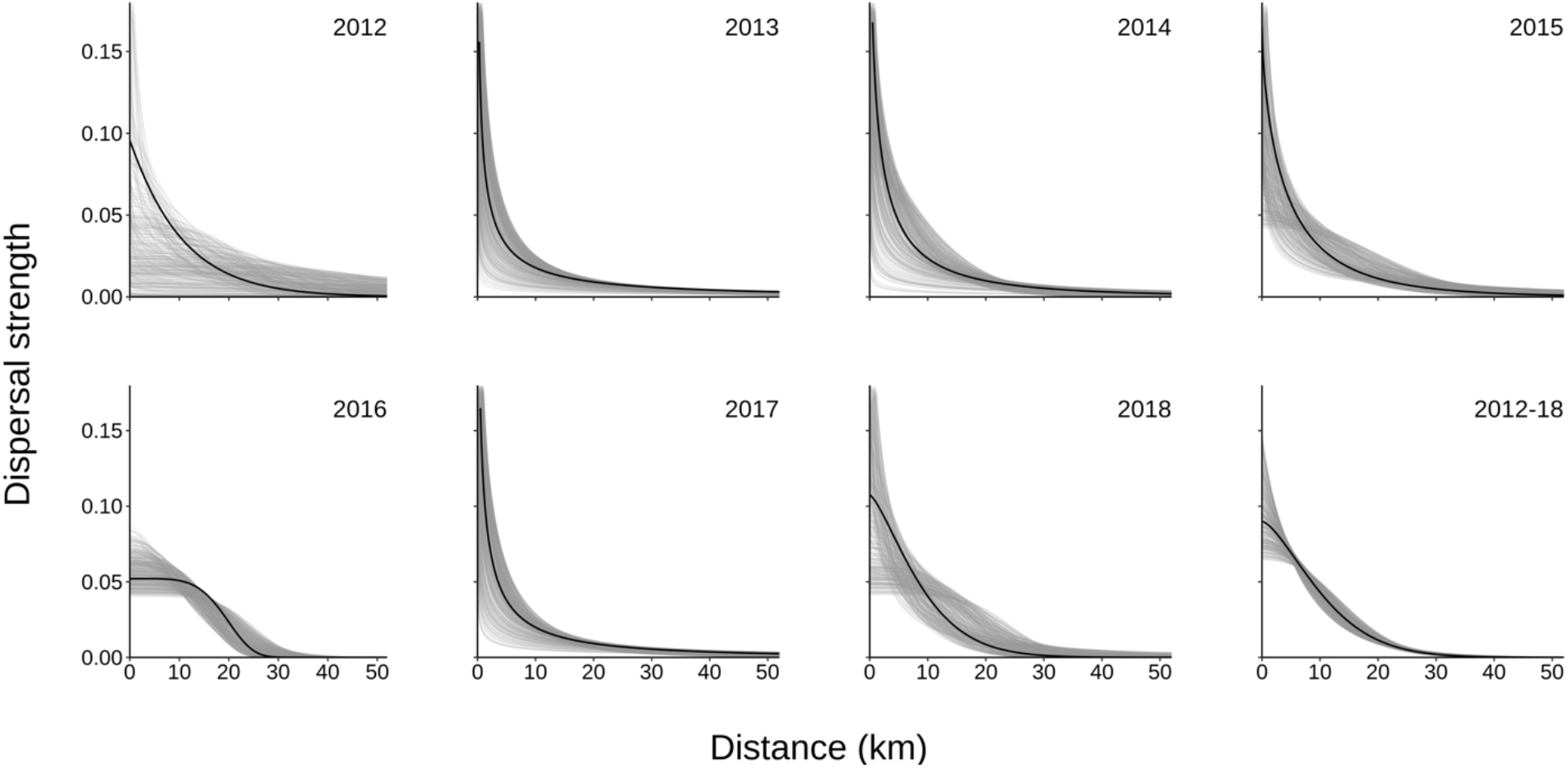
Annual fitted dispersal kernels describing the strength of dispersal given distance for each individual year and for all years combined. Gray lines behind the point estimate of the kernel (black) are 500 kernels drawn from the 95% CI likelihood surface.

Mean dispersal distance estimates had wide confidence bounds but were concentrated at or below 10 km in 2012, 2015, 2016, 2018, and the average kernel, and were spread from 11 to 70 km in 2013, 2014, and 2017 (Fig. 4A, Table S2). Mean dispersal distance point estimates differed almost ten-fold across years (CV = 1.03), and these differences were greater than expected from the null model (*p* = 0.001, Fig. S3C).

**Figure 4.**
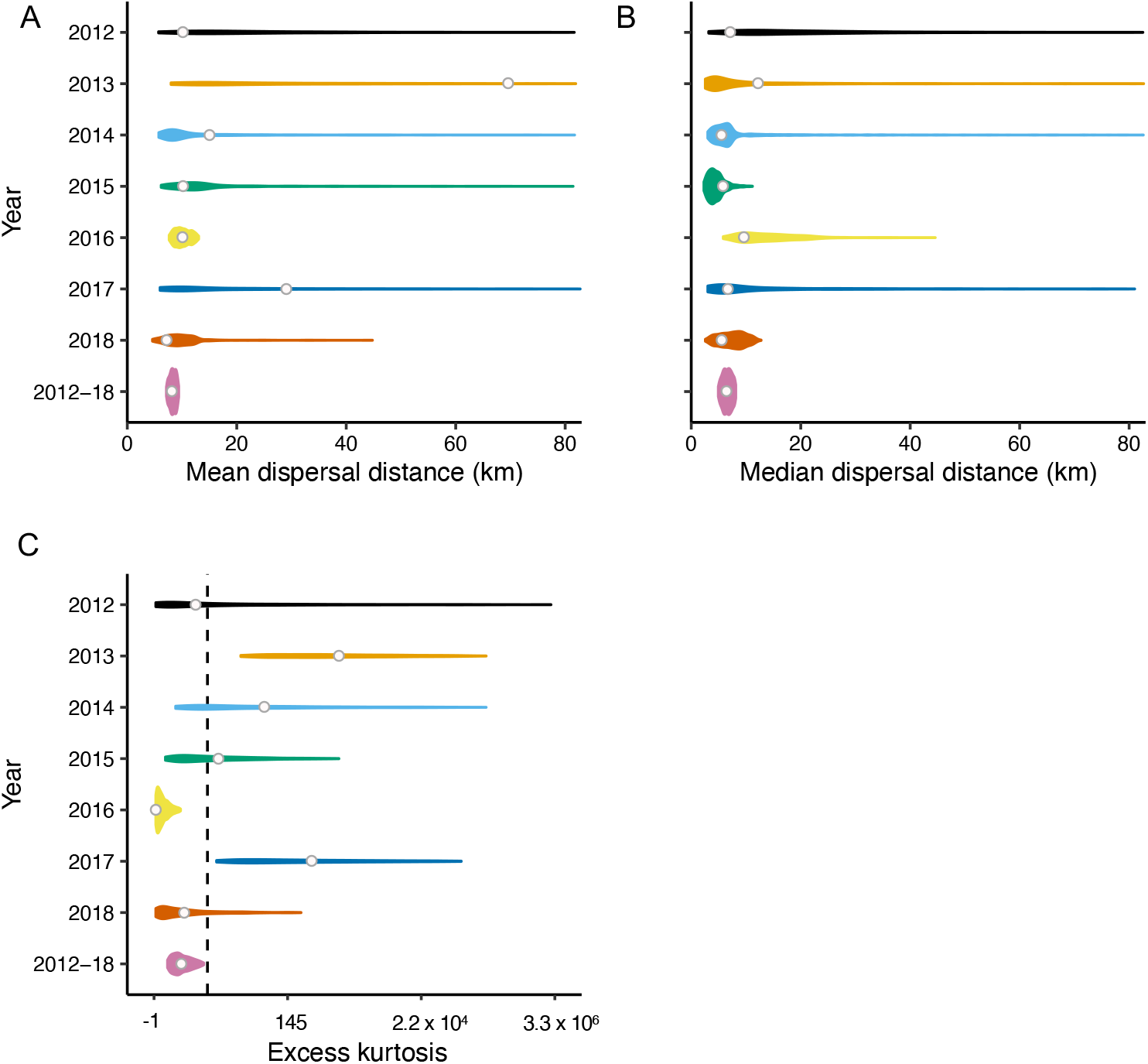
Point estimates (white circle) and uncertainty distributions of the A) mean dispersal distance, B) median dispersal distance, and C) excess kurtosis for each annual kernel and for all years combined. Excess kurtosis less than 2 (black vertical line) indicates a platykurtic (thin-tailed) and positive values a leptokurtic (fat-tailed) distribution relative to a Gaussian kernel.

Median dispersal distances ranged from a minimum of 5 km in 2013 to 12 km in 2012 (CV = 0.34), corresponding to a two-fold difference among years (Fig. 4B). In every year, 50% of larvae recruited within 15 km or less of their natal origin (Fig. 4B, Table S2) but the variation among years was greater than expected from the null model (*p* = 0.041, Fig. S3D). The distance at which 90% of larvae recruited ranged from a minimum of 15 km to a maximum of 157 km, while the proportion of recruits retained in the 30 km study site was between 0.33-0.49 across years (Table S2).

Driven by differences in kernel shape, we observed variation in the excess kurtosis among kernels (Fig. 4C). The best-fits for 2012, 2016, 2018, and the average kernel were thin-tailed. In contrast, the best-fits for 2013, 2014, 2015, and 2017 were fat-tailed and suggested a greater proportion of long-distance dispersal events.

The dispersal kernels fit to Northeast Monsoon and Southwest Monsoon recruits differed in scale (SD = 1.21) and shape (CV = 0.67) (Fig. 5, Fig. S4 A&B, Table S3), and both were significantly greater than the null model (*p* = 0.001 for scale and shape). Mean dispersal distance was 8.6 km for the Northeast Monsoon and 9.5 km for the Southwest Monsoon, and this small difference (CV = 0.07) was not significant (*p* = 0.319 when compared to the null model, Fig. S4C). Median dispersal distance was greater in the Southwest Monsoon (6.9 km) than in the Northeast Monsoon (4.8 km) and this difference was significant (CV=0.25, *p* = 0.003 from the null model, Fig. S4D, Table S3). The estimated Northeast Monsoon kernel was fat-tailed (θ = 0.56 [0.54, 0.61]), while the Southwest Monsoon kernel was less fat-tailed (θ = 1.58 [1.34, 1.59]), and so the distance at which 90% of larvae recruited was greater in the Northeast than the Southwest Monsoon (23.6 vs 18.3). The proportion of recruits retained within the metapopulation was 0.47 in the Northeast Monsoon and 0.49 in the Southwest Monsoon (Table S3).

**Figure 5.**
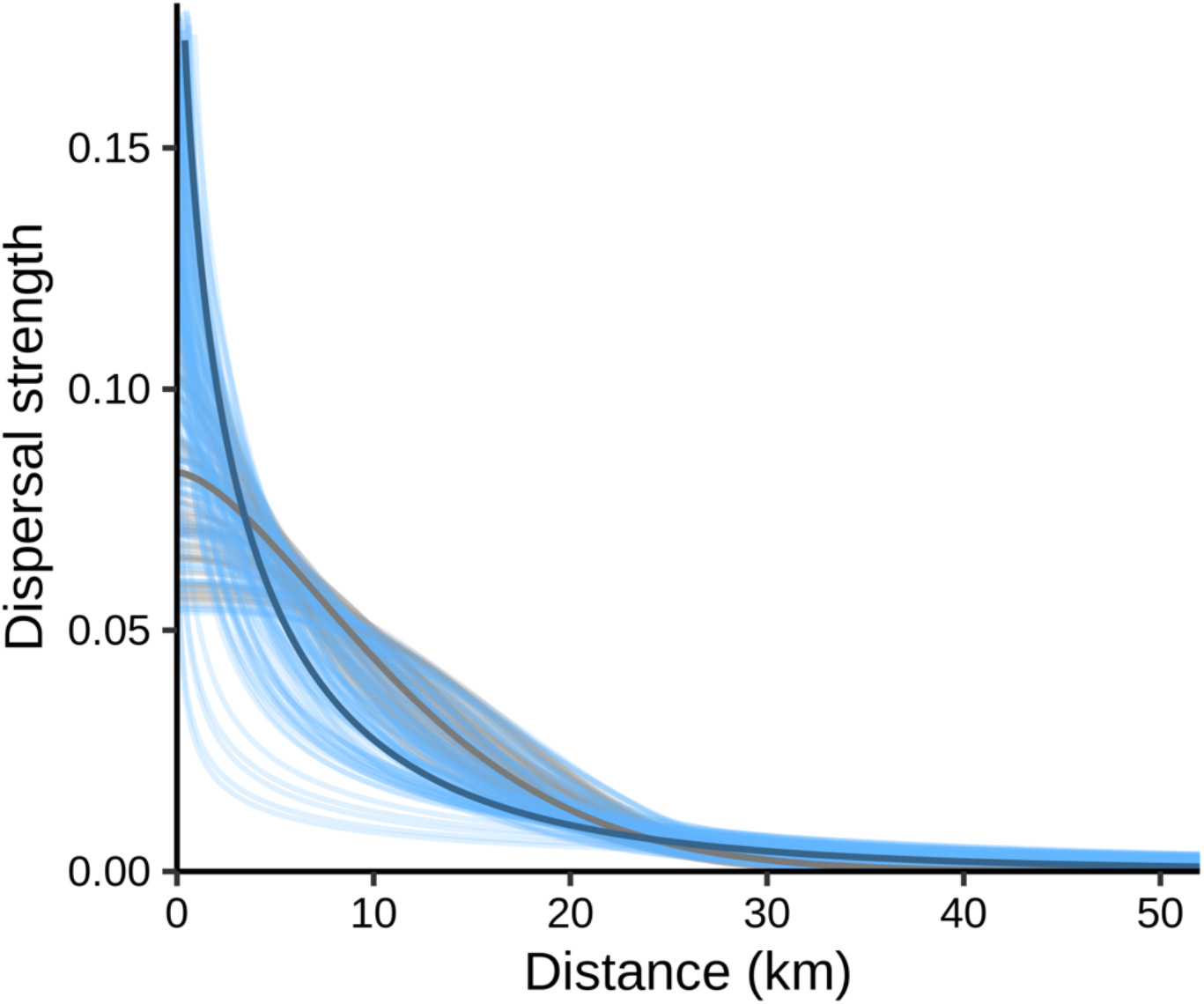
The seasonal dispersal kernels for recruits likely dispersing in the Northeast Monsoon season (blue) or the Southwest Monsoon season (gray). Lines behind the point estimate of the kernel are 500 kernels drawn from the 95% CI likelihood surface.

Fitting kernels to down-sampled datasets demonstrated that *k* and θ were relatively insensitive to samples larger than 50% of our dataset, but that *k* and θ were biased high for smaller datasets (Fig. S5, Fig. S6).

### Variation in dispersal direction

The most parsimonious regression model suggested that the probability of dispersal between any two patches declined with distance, was higher for dispersal from north to south, increased with the size of destination and source habitat patches, and varied among years (Table S5). There was some evidence for including recruit sampling effort (ΔAIC = 1.8) as well as adult sampling effort (ΔAIC = 2). The interaction between year and direction was not included in any of the models with ΔAIC < 5, suggesting no evidence for change in the predominant direction of dispersal among years (Table S5).

The best model for the seasonal kernels was similar, but there was some evidence for variation in dispersal direction among seasons (Table S6). A model with ΔAIC = 2 included the direction by year interaction term. The best model suggested predominantly southwards dispersal in the Northeast Monsoon season (wind blowing out of the northeast) and predominantly northwards dispersal in the Southwest Monsoon season (wind blowing out of the southwest, Fig. 2, Table S6B).

### Correlation among dispersal routes

Self-recruitment to sites was low, from a minimum of 0.23% to a maximum of 8.6%. However, dispersal probabilities along individual routes had high interannual coefficients of variation that were generally highest for self-recruitment routes (range = 0.45-1.0, mean = 0.53, Fig. 6A). Temporal variation in self-recruitment rates for the smallest sites was particularly high, as was variation in routes connecting the largest sites (Fig. 6A). For example, self-recruitment in the smallest site (Poroc San Flower) was estimated to be eight times larger in 2017 (1.8%) than in 2016 (0.23%).

**Figure 6.**
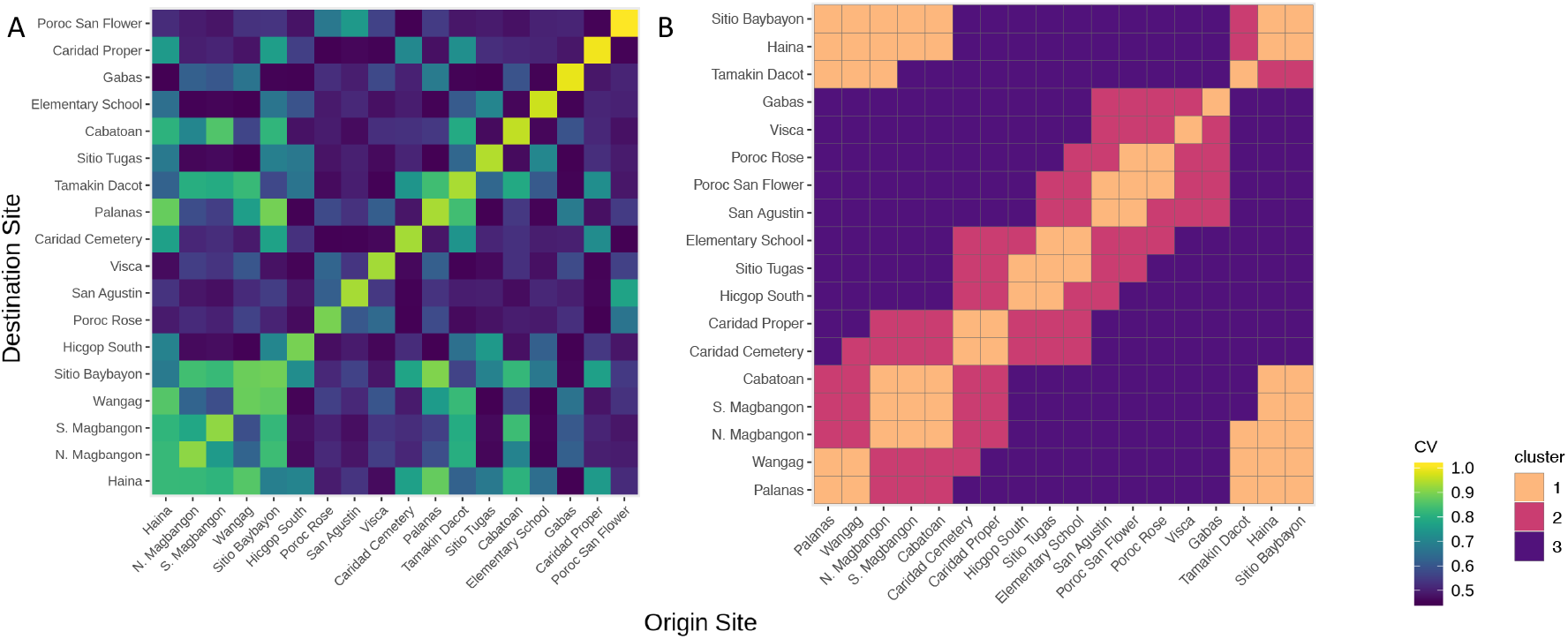
A) Coefficient of variation (CV) for each dispersal route across seven years (2012-2018). Sites are ordered by size from the bottom-left (Haina is the largest and Poroc San Flower the smallest). B) Cluster groupings identified from the correlation matrix among routes, arranged from the northernmost (Palanas) to southernmost (Sitio Baybayon) site.

We also found strong correlations between dispersal routes. Coefficients ranged from −0.94 to 1, with a mean correlation of 0.21 ± 0.67 (SD) (Fig. 6B). Clustering identified three main groups (Fig. S7). The first cluster included routes between sites that were both geographically close (including self-recruitment) and distant (−0.28 ± 0.61). Within this cluster, routes between close sites were positively correlated, and distant routes were negatively correlated. The second cluster included routes between sites that were somewhat close to each other and were more weakly correlated (0.23 ± 0.53). The third group was of routes between sites that were intermediately distant and mostly positively correlated (0.36 ± 0.66).

## Discussion

Quantifying variation in metapopulation connectivity across time remains an important challenge given the implications for metapopulations, metacommunities, and evolutionary processes (Bode et al., 2017; G. P. Jones et al., 2009; Levin, 2006). Here, we directly quantified dispersal patterns across seven years in a coral reef fish and found substantial variation in dispersal kernel scale and shape across years. We also observed seasonal variation in dispersal that was of similar magnitude to interannual variation. Though there was temporal variation in dispersal distance, we found some evidence of variation in direction between monsoon seasons but did not find strong evidence for variation in the dispersal direction among years. Self-recruitment at the scale of individual sites and of the metapopulation varied strongly through time, and dispersal probabilities between sites were often correlated.

Marine larval dispersal has been called the “black box” of marine ecology (Robert K. Cowen, 2002), and our results contribute to the growing understanding of differences among species. Mean dispersal distances for the congener *Amphiprion percula* (10-15 km) are similar to the average (8 km) we found for *A. clarkii* (Almany et al., 2017). However, mean dispersal distances in other coral reef fishes range from 2.8 km in neon gobies (D’Aloia et al., 2015) to 64 km or greater in groupers and butterflyfishes (Almany et al., 2017; Williamson et al., 2016). A focus on average patterns, however, misses the fact that marine dispersal patterns vary strongly through time. Indeed, the interannual differences in mean (7 to 70 km) dispersal distance that we observed are comparable in magnitude to the differences across reef fish species (Almany et al., 2017; Bode et al., 2017; D’Aloia et al., 2015; Williamson et al., 2016). Previous studies have clarified with simulations that such temporal variation can occur (Bani et al., 2019; Snyder et al., 2014; James R. Watson et al., 2012), and that some variation among two to three years does occur empirically (Almany et al., 2017; Berumen et al., 2012; Hogan et al., 2012). However, the degree and character of dispersal variation has not been clear. Across seven years, we found that kernel scale, shape, the resulting mean dispersal distances varied substantially.

The variation in the proportion of recruits retained in the metapopulation among years suggests differing degrees of larval retention within the sampled metapopulation. Larval retention is a key metric because these recruits can replace individuals lost to mortality and thereby allow the metapopulation to persist (Burgess et al., 2014). In years with thin-tailed kernels, our estimates suggest most larvae were retained within the metapopulation (up to 50%), while years with fat-tailed kernels likely dispersed substantial proportions of larvae beyond the metapopulation with 90% of larvae recruiting at distances as large as 160 km in 2013. While converting these retention fractions to persistence requires information on lifetime egg production, rough calculations suggest that retention of 31-35% could be enough for persistence (Botsford et al., 2009; Burgess et al., 2014), implying that our sampled metapopulation may reach persistence criteria in most years. The cross-year average kernel was approximately Laplacian in shape and yet kernels in some years were strongly thin-tailed or fat-tailed, suggesting that an average measure of dispersal does not reflect the importance of rare, long-distance dispersal events in certain years. We do note, however, that our observations were limited to distances within 30 km and so did not sample the tails directly. These results caution against considering only one estimate of dispersal to understand ecological or evolutionary dynamics or for making conservation decisions, especially if dispersal is estimated from a single year of observations.

Additionally, we observed that dispersal estimates varied across monsoon seasons with a similar magnitude to variation among years. Previous work has demonstrated that the predominant direction of dispersal in marine mussel species can differ seasonally, driven by alternating wind-driven surface currents (Carson et al., 2010), and that reef fish species are similarly affected by seasonal currents (Abesamis *et al.* 2017). Here, we observed differences in the scale and shape of the dispersal kernel between seasons, with limited evidence for differences in directionality. Our results suggest somewhat greater retention of larvae within the metapopulation during the Southwest Monsoon season and greater dispersal beyond the metapopulation in the Northeast Monsoon season. The Southwest Monsoon brings a period of heavy rainfall and westerly winds, coinciding with the tropical cyclone season, while the Northeast Monsoon is a dry season dominated by easterly winds (Lau & Yang, 1997). The onset of the Southwest Monsoon is variable on both annual and decadal time scales (Kubota, Shirooka, Matsumoto, Cayanan, & Hilario, 2017; Lau & Yang, 1997). This seasonal shift in the atmosphere is likely to alter oceanographic flow and therefore influence larval dispersal patterns (Robert K Cowen & Sponaugle, 2009).

Alternatively, the seasonal variation we observed could result from seasonal differences in recruit survival rather than differences in dispersal. For example, larvae of *Stegastes partitus* grow faster and recruit at smaller sizes in warmer water, and these smaller larvae also experience greater mortality than their larger conspecifics that develop in cooler water, although this pattern can reverse seasonally (Rankin & Sponaugle, 2011). The Southwest Monsoon recruits we sampled had to survive from settlement until sampling and so were approximately six months older than recruits from the Northeast Monsoon. Rather than temporal variation in dispersal patterns, therefore, the seasonal differences in dispersal kernels could indicate post-settlement selection against long-distance dispersers, as has been suggested from phenotype-environment mismatches (Marshall, Monro, Bode, Keough, & Swearer, 2010).

While we studied a marine system, seasonal variation in dispersal is also likely in terrestrial systems. In deciduous forests, the seasonal loss of foliage can increase the potential for long-distance dispersal by allowing vertical eddy motions to transport seeds higher in the canopy where winds are stronger (R. Nathan & Katul, 2005). Broadly, seasonal variation in reproduction should increase variability in connectivity patterns, not only because environments can differ between seasons (e.g., monsoons or storm frequency), but also because seasonal reproduction will sample a smaller range of environmental conditions and therefore be more sensitive to chaotic or stochastic environmental variation (Siegel et al., 2008; Snell et al., 2019; Snyder et al., 2014).

Variation in connectivity likely drives interannual differences in the contribution of the metapopulation to its own growth and persistence. When combined with demographic rates such as survival and lifetime reproductive output, the kernels we estimate here can be used to estimate the metapopulation growth rate and thus persistence (Burgess et al., 2014; Hastings & Botsford, 2006). In simulations, variability in dispersal often diminishes but sometimes enhances the persistence of metapopulations (Snyder et al., 2014; James R. Watson et al., 2012; Williams & Hastings, 2013), though empirical observations to test these predictions have been lacking. The impact of variable connectivity on metapopulation growth rates is expected to depend largely on how dispersal routes are correlated. When connections are positively correlated, stochastic growth rates are predicted to be lower than growth rates calculated using a time-averaged estimate of connectivity (Snyder et al., 2014; James R. Watson et al., 2012). However, dispersal variability can instead enhance metapopulation persistence if dispersal routes are anti-correlated, such as with complementary dispersal regimes caused by current flows that alternate the directionality of dispersal (Williams & Hastings, 2013). Our regression model indicated that the average direction of dispersal does not vary substantially across years, but we did observe both negative and positive correlations between dispersal routes. Intuitively, connections that were geographically close were positively correlated, while distant connections could also be negatively correlated. Overall, dispersal routes in this metapopulation were positively correlated, suggesting that stochastic metapopulation growth rates are likely lower than those expected from time-averaged connectivity.

Our observations of variation in dispersal are the most comprehensive, empirical time-series to date, and they demonstrate variability on interannual and seasonal time scales in a metapopulation as predicted by biophysical simulations of connectivity (James, Armsworth, Mason, & Bode, 2002; Mitarai et al., 2008; Siegel et al., 2008; Snyder et al., 2014). Biophysical simulations of dispersal in fluid environments (atmosphere and ocean) are a promising avenue for predictive models of dispersal that can capture environmental stochasticity in connectivity across time for diverse organisms such as plants, marine larvae, insects, and fungi (Lett et al., 2020). As an important step towards synthesizing biophysical simulations and empirical dispersal observations, recent work has demonstrated that biophysical simulations coupled with species-specific behaviors can explain dispersal patterns observed directly through parentage matches (Bode et al., 2019). An important next step will be to understand whether such biophysical simulations can also explain temporal variability in dispersal kernels. Such an approach would provide insight into the environmental mechanisms contributing to variation in connectivity patterns.

Although these results are the most complete time series of dispersal kernels of which we are aware and cover more than a generation in our focal species, seven years remains limited. Future work assessing longer time series, ideally on decadal scales that may illustrate the importance of long-term climatic oscillations, is an important though logistically challenging next step. There is also a need for studies of dispersal variability across diverse environments and taxa to assess the generality of these results. Additionally, we described dispersal using dispersal kernels because they provide a convenient framework for considering dispersal across spatial scales. However, kernels are isotropic and assume the functional form of the true underlying distribution, which makes it difficult to make inferences about the mechanisms driving dispersal patterns (Bode et al., 2019) or to assess details about site-to-site dispersal. Future empirical work focusing on a mechanistic understanding of dispersal variability and its ecological and evolutionary impacts will illuminate both the ecological and evolutionary dynamics of metapopulations.

The variation in dispersal kernels observed here likely occurs in many other organisms governed by metapopulation dynamics in the nearshore environment (Mitarai et al., 2008; Siegel et al., 2008), as well as in terrestrial organisms where dispersal is subject to heterogenous environmental forcings such as wind-driven seed dispersal in plants (R. Nathan & Katul, 2005; Ran Nathan et al., 2008; Snell et al., 2019), spore dispersal in fungi (Geml, Tulloss, Laursen, Sazanova, & Taylor, 2008), or silk ballooning dispersal in arthropods (Bell, Bohan, Shaw, & Weyman, 2005). Stochastic connectivity will be especially important for organisms with seasonal reproduction and structurally complex dispersal landscapes (Siegel et al., 2008). While our understanding of time-averaged patterns of connectivity across diverse taxa and systems has improved substantially, describing the variation around the mean and the consequences of such variation is a much-needed next step.

## Acknowledgements

For financial support, we thank the Rutgers School of Environmental and Biological Sciences, the Rutgers Institute of Earth, Ocean, and Atmospheric Sciences, the US National Science Foundation (#OCE-1430218 and #OCE-1426891), an Alfred P. Sloan Research Fellowship, and an Oak Ridge Associated Universities Ralph E. Powe Junior Faculty Enhancement award. We thank both Michael Bode and Lauren Sullivan for their comments on a draft manuscript. We thank Jennifer Hoey, Patrick Flanagan, Joyce Ong, Geralde Sucano, Beverlito Montalban, Cecil Bantiles, Teresita Idara, Rogello Nicanor, Liza Espinosa, Shem San Jose, Noel Alquino, Carlos Balansuna, Froilan Beñas, Danilo Marine, Tony Nahacky, Apollo Lizano, Rodney Silvado, Arturo Bastasa, the municipalities of Bay Bay City and Albuera, and Visayas State University for their support of our field sampling in Leyte, Philippines. We also thank Dave Portnoy and John Gold for assistance with ddRADseq lab protocols.

## Data accessibility statement

The population genomic data, myBaits probe sequences, sample metadata, GIS data, and all accompanying bioinformatics and R analysis code to produce these results is publicly available as a GitHub repository (https://github.com/pinskylab/DispersalVariation) and will be archived with a DOI through Zenodo. The population genomic data will also be archived the NCBI Sequence Read Archive.

## Statement of authorship

KAC and MLP conceived of and designed the study. MRS, MLP, KAC, and AGD collected field data. HRM facilitated field work. MRS led molecular wet lab work. MRS and JBP conducted bioinformatics analysis. KAC analyzed genotype data and wrote the manuscript. All authors contributed to revisions.

