## Supporting Information for "Quantifying dispersal variability among nearshore marine populations"

***Field sampling***

Sites were sampled in late May and June, except for 2018 when they were sampled in March and April (Table S1). We captured fish with drive nets and dip nets while on SCUBA and measured their fork length. For fish with a fork length greater than or equal to 3.5 cm, we non-lethally sampled a portion of the caudal fin and preserved it in 95% ethanol. In 2017, we also collected recruits less than 3.5 cm. We tagged the host anemone with a unique identification number and recorded the location using GPS coordinates. We categorized recruits as less than 6.0 cm and adults as greater than or equal to 6.0 cm based on the size distributions of known subadults and breeders and estimates of annual growth from mark recapture studies in this population.

***DNA sequencing and bioinformatics***

We extracted genomic DNA from tissue samples using DNeasy Blood & Tissue Kits (Qiagen; Hilden, Germany) according to manufacturer protocols, evaluated the extractions on 2% agarose gel, and quantified them using PicoGreen and a SpectraMax M3 Microplate Reader (Thermo Fisher Scientific, Waltham, MA, USA; Molecular Devices, Sunnyvale, CA, USA). Following a ddRADseq protocol for genomic library preparation (Peterson *et al.* 2012), we digested the DNA with PstI and EcoRI restriction enzymes, then ligated one of 48 custom barcodes to the DNA fragments by individual (Table S7). Next, we pooled DNA into groups of 48 individuals and size selected to 375 +/- 37 base pairs (bp) using a Sage Science Pippin Prep (Sage Science, Beverly, MA). We added one of 12 indexed Illumina primers to the pooled DNA fragments using polymerase chain reaction (PCR) (Table S8). PCR products of samples from field seasons 2012-2015 were pooled again into final DNA sequencing libraries containing 192 individuals. PCR products of samples from field seasons 2016-2018 underwent in-solution sequence capture with a myBaits kit following the Arbor Bioscience protocol (Arbor Bioscience, Ann Arbor, MI, USA) and pooled into libraries containing 576 individuals. Libraries were sequenced on an Illumina 2500 (single end, 150 bp) at the Princeton Core Facility (Princeton, NJ, USA) (Table S9).

We used process-radtags (Catchen *et al.* 2013) to demultiplex the raw sequencing data and then used the bioinformatics pipeline dDocent to trim and map adaptor sequences to a *de novo* ddRAD contig assembly (Puritz *et al.* 2014). Next, we used the program FreeBayes (Garrison & Marth 2012) to call SNPs. To isolate SNPs most useful for parentage assignment, we used PLINK 1.9 (Purcell *et al.* 2007) to remove loci not in Hardy Weinberg equilibrium and kept only a single SNP of those in linkage disequilibrium (r^2^ cutoff of 0.5). We then filtered SNPs using VCFTools (Danecek *et al.* 2011) for a minor allele frequency of at least 0.2 and a minimum sequencing depth of 16x across all individuals. We then removed individuals with a genotyping rate less than 70% across all remaining loci (O’Leary *et al.* 2018). Finally, we removed indels and converted 1,340 SNPs on 1,050 contigs to a Genepop file for analysis.

***Fitting dispersal kernels***

We used R 3.10.0 (R Core Team, 2019) and the package bbmle v1.0.20 (Bolker & Team 2017) to search for the maximum likelihood estimates of *k* (from -10 to 10) and θ (from 0.1 to 5) using the Limited-memory BFGS algorithm and the dispersal kernel likelihood function from Bode et al. (2017). In the kernel fitting procedure, both assigned and unassigned recruits are informative because unassigned recruits indicate distances to which there is a low probability of dispersal from sampled parents (Bode *et al.* 2017). The probability of finding a parentage match also depends on the proportion of total individuals sampled; as more individuals are sampled, the probability of finding a match increases. The likelihood function uses the proportion of individuals sampled at each site to assign expected recruits to sources and scales the expected recruits according to the estimated site size. We used habitat to calculate the proportion of each site sampled by dividing the number of anemones visited in a sampling year by the total number of anemones ever observed at a given site. Because anemones experience mortality, this calculation may therefore overestimate total anemones and underestimate the proportion of habitat that we sampled. However, the accuracy of the likelihood method is not markedly improved beyond sampling 10% of the adult population (Bode *et al.* 2017), and the site specific estimates of proportion sampled were rarely lower than 0.1 (0.49 $\pm$ 0.31, mean $\pm$SD). Additionally, from mark-recapture studies in Ormoc Bay, we estimate annual anemone survival to average 75%, and so the potential bias is expected to be relatively small.

The kernel fitting method accounts for unassigned recruits at each patch (Bode et al. 2017). The function calculates the likelihood that these recruits came from either an unsampled parent on a sampled patch or an unsampled “ghost” population that represents all sites outside of our focal region in Ormoc Bay. To represent “ghost” populations, we identified other unsampled reef patches in the Camotes Sea from previous survey work in the region (Pinsky *et al.* 2010). We included the nearshore areas of the Camotes Islands and Cuatros Islas as well as the Leyte coastline 10 km north and 10 km south of our focal region. For each of these locations, we assumed coral reef habitat to be within 0.5 km of the coastline and estimated the area of these “ghost” populations using the GIS software QGIS v2.18. We estimated that 10% percent of the area of these locations were habitable reef, a conservative estimate based on the reef cover of our focal region.

**Table S1 Genotyped individuals included in each annual analysis**

Summary of all genotyped recruits and adults that were included in each annual analysis and of the total number of genotyped recruits and adults across the entire study (2012-2018). We cumulatively pooled adults across years so that once an adult was sampled, it was included as a candidate parent in the analyses of that and all subsequent years.

| Analysis year | Recruits | Adults |
| --- | --- | --- |
| 2012 | 63 | 157 |
| 2013 | 150 | 372 |
| 2014 | 181 | 536 |
| 2015 | 111 | 850 |
| 2016 | 111 | 1,232 |
| 2017 | 130 | 1,453 |
| 2018 | 45 | 1,729 |
| Total | 791 | 1,729 |

**Table S2** **Parentage assignments and kernel fits**

| Year | Assignment percentage | *k*  [95% CI] | θ  [95% CI] | Mean dispersal distance (km)  [95% CI] | Median dispersal distance (km)  [95% CI] | Distance (km)  90% dispersal | Proportion retained in 30 km |
| --- | --- | --- | --- | --- | --- | --- | --- |
| 2012 | 4.8 % | -2.36 [-7.22, -1.9] | 1.03 [0.48, 1.10] | 10.2 [0, 250+] | 7.1 [3.2 - 250+] | 23.3 | 0.47 |
| 2013 | 14.0 % | 4.04 [3.59, 4.06] | 0.22 [0.20, 0.24] | 69.6 [8.0, 250+] | 12.2 [2.6- 250+] | 157.4 | 0.33 |
| 2014 | 7.2 % | 0.49 [-0.26, 0.81] | 0.38 [0.37, 0.4] | 15.0 [5.6, 250+] | 5.5 [3.0- 250+] | 38.1 | 0.43 |
| 2015 | 9.9 % | -1.52 [-2.08, -1.13] | 0.67 [0.56, 0.70] | 10.2 [6.1, 240.5] | 5.8 [2.3- 11.2] | 24.9 | 0.47 |
| 2016 | 5.4 % | -3.04 [-3.07, -2.88] | 5 [2.16, NA] | 10.1 [7.6, 13.1] | 9.6 [5.9- 44.6] | 18.9 | 0.50 |
| 2017 | 10 % | 2.94 [2.66, 3.64] | 0.26 [0.23, 0.26] | 29.1 [5.9, 250+] | 6.7 [3.0- 250+] | 69.7 | 0.39 |
| 2018 | 8.9 % | -2.32 [-2.51, -2.3] | 1.37 [1.36, 1.60] | 7.2 [4.5, 50.8] | 5.6 [2.6- 12.8] | 15.7 | 0.50 |
| 2012-  2018 | 9.0 % | -2.51 [-2.51, -2.48] | 1.49 [1.32, 1.60 | 8.2 [7.1, 9.4] | 6.4 [4.9- 8.1] | 17.6 | 0.50 |

Parentage assignment percentage, maximum likelihood estimates (MLE) of the dispersal kernel parameters, mean and median dispersal distances of the resulting kernels, the distance at which 90% of larvae recruit, and estimated proportion of recruits retained within the 30 km study system for each individual year and for all years combined (2012-2018). Parameter *k* defines the scale of the distribution and parameter θ defines the shape of the distribution. Confidence intervals (CI) are the 95% likelihood surface values of *k* and θ. When median or mean dispersal distance was calculated to be greater than 250 km because the dispersal kernel was fat-tailed, we indicated that as 250+. For 2016, the upper bound of θ was undefined and is marked NA.

**Table S3 Seasonal kernel fits**

Parentage analysis assignment percentage for sampled recruits of the Northeast Monsoon and Southwest Monsoon seasons, maximum likelihood estimates (MLE) of the parameter sets describing the dispersal kernel, the mean and median dispersal distances of the resulting kernels, the distance at which 90% of larvae recruit, and the estimated proportion of recruits retained within the 30 km study system. Parameter *k* defines the scale of the distribution and parameter θ defines the shape of the distribution. Confidence intervals (CI) are the 95% likelihood surface values of *k* and θ. When dispersal distance was calculated to be greater than 250 km because the dispersal kernel was fat-tailed, we indicated that as 250+.

| Season | Assignment percentage | *k*  [95% CI] | θ  [95% CI] | Mean dispersal distance (km)  [95% CI] | Median dispersal distance (km)  [95% CI] | Distance (km)  90% dispersal | Proportion retained in 30 km |
| --- | --- | --- | --- | --- | --- | --- | --- |
| Northeast Monsoon | 1.4% | -0.89 [-1.46, -0.74] | 0.56 [0.54, 0.61] | 9.5 [6.0, 250+] | 4.8 [3.2, 250+] | 23.6 | 0.47 |
| Southwest Monsoon | 4.4% | -2.60 [-2.63, -2.47] | 1.58 [1.34, 1.59] | 8.6 [7.1, 10.4] | 6.9 [4.9, 9.3] | 18.3 | 0.49 |

**Table S4 Annual kernel shape differences**

Differences in kernel shapes based on the CI overlap of the MLE of θ. If the CIs did not overlap between two fits, we marked the fits as different. An asterisk (*) denotes a different kernel shape and a hyphen (-) denotes no difference.

| Year | 2013 | 2014 | 2015 | 2016 | 2017 | 2018 | 2012-  2018 |
| --- | --- | --- | --- | --- | --- | --- | --- |
| 2012 | * | - | - | * | * | * | * |
| 2013 |  | * | * | * | * | * | * |
| 2014 |  |  | * | * | * | * | * |
| 2015 |  |  |  | * | * | * | * |
| 2016 |  |  |  |  | * | * | * |
| 2017 |  |  |  |  |  | * | * |
| 2018 |  |  |  |  |  |  | - |

**Table S5 Annual dispersal direction model results**

Results of the logistic regression explaining whether or not a dispersal event was observed along each possible dispersal route. Possible directions of dispersal events were North or South (relative to parent patch) and self (self-recruitment). A) AIC results used in model selection. The best-fit model is bolded. B) Coefficients for the best-fit model on a logit scale.

A) Model selection

| Models compared | d.f. | AIC | Residual Df | Residual Dev | ΔAIC | ΔAIC Weight |
| --- | --- | --- | --- | --- | --- | --- |
| Distance + Source habitat size + Destination habitat size + Direction + Year | 11 | 411.76 | 2516 | 389.76 | 0 | 0.49 |
| Distance + Adult sampling effort + Source habitat size + Destination habitat size + Direction + Year | 12 | 413.54 | 2515 | 389.54 | 1.8 | 0.2 |
| Distance + Recruit sampling effort + Source habitat size + Destination habitat size + Direction + Year | 12 | 413.74 | 2515 | 389.74 | 2 | 0.18 |
| Distance + Recruit sampling effort + Adult sampling effort + Source habitat size + Destination habitat size + Direction + Year | 13 | 415.51 | 2514 | 389.51 | 3.8 | 0.08 |
| Distance + Source habitat size + Destination habitat size + Direction * Year | 17 | 416.82 | 2510 | 382.82 | 5.1 | 0.04 |
| Distance + Recruit sampling effort + Adult sampling effort + Source habitat size + Destination habitat size + Direction * Year | 19 | 420.63 | 2508 | 382.63 | 8.9 | 0.01 |
| Distance + Recruit sampling effort + Adult sampling effort + Destination habitat size + Direction + Year | 12 | 446.48 | 2515 | 422.48 | 34.7 | 0 |
| Distance + Recruit sampling effort + Adult sampling effort + Source habitat size + Direction + Year | 12 | 450.27 | 2515 | 426.27 | 38.5 | 0 |
| Distance + Recruit sampling effort + Adult sampling effort + Source habitat size + Destination habitat size + Direction | 5 | 483.69 | 2522 | 473.69 | 71.9 | 0 |
| Distance | 2 | 485.26 | 2525 | 481.26 | 73.5 | 0 |

B) Summary of best-fit model

| Parameter | Estimate | Standard Error | z-statistic | P value |
| --- | --- | --- | --- | --- |
| (Intercept) | -4.598 | 0.598 | -7.693 | 1.4x10^-14^ |
| Distance (km) | -0.052 | 0.018 | -2.884 | 0.004 |
| Direction (-1, 0, 1) | -0.160 | 0.164 | -0.973 | 0.330 |
| Destination habitat size | 0.002 | 0.000 | 6.143 | 8.11 x10^-10^ |
| Source habitat size | 0.002 | 0.000 | 6.306 | \| 2.87 x10^-10^ |
| Year 2013 | 1.098 | 0.654 | 1.679 | 0.093 |
| Year 2014 | 0.261 | 0.698 | 0.373 | 0.709 |
| Year 2015 | 0.011 | 0.707 | 0.015 | 0.988 |
| Year 2016 | -1.376 | 0.837 | -1.645 | 0.100 |
| Year 2017 | -1.466 | 0.839 | -1.746 | 0.081 |
| Year 2018 | -3.493 | 1.104 | -3.164 | 0.002 |

**Table S6 Seasonal dispersal direction model results**

Results of the logistic regression explaining the dispersal events observed from the Northeast Monsoon season and Southwest Monsoon season for all possible dispersal routes. Possible directions of dispersal events were North or South (relative to parent patch) and self (self-recruitment). A) AIC results used in model selection. The best-fit model is bolded. B) Model coefficients for the best-fit model.

A) Model selection

| Models compared | d.f. | AIC | Residual. Df | Residual. Dev | ΔAIC | ΔAIC Weight |
| --- | --- | --- | --- | --- | --- | --- |
| Distance + Source habitat size + Destination habitat size + Direction + Season | 6 | 197.22 | 716 | 185.22 | 0 | 0.4 |
| Distance + Source habitat size + Destination habitat size + Direction * Season | 7 | 198.76 | 715 | 184.76 | 1.5 | 0.18 |
| Distance + Recruit sampling effort + Source habitat size + Destination habitat size + Direction + Season | 7 | 198.88 | 715 | 184.88 | 1.7 | 0.17 |
| Distance + Adult sampling effort + Source habitat size + Destination habitat size + Direction + Season | 7 | 199.22 | 715 | 185.22 | 2 | 0.15 |
| Distance + Recruit sampling effort + Adult sampling effort + Source habitat size + Destination habitat size + Direction + Season | 8 | 200.88 | 714 | 184.88 | 3.7 | 0.06 |
| Distance + Recruit sampling effort + Adult sampling effort + Source habitat size + Destination habitat size + Direction * Season | 9 | 202.43 | 713 | 184.43 | 5.2 | 0.03 |
| Distance + Recruit sampling effort + Adult sampling effort + Source habitat size + Direction + Season | 7 | 223.40 | 715 | 209.40 | 26.2 | 0 |
| Distance + Recruit sampling effort + Adult sampling effort + Source habitat size + Direction + Season | 7 | 242.66 | 715 | 228.66 | 45.4 | 0 |
| Distance | 2 | 271.97 | 720 | 267.97 | 74.7 | 0 |
| Distance + Recruit sampling effort + Adult sampling effort + Source habitat size + Destination habitat size + Direction | 5 | 273.31 | 717 | 263.31 | 76.1 | 0 |

B) Summary of best-fit model

| Parameter | Estimate | Standard Error | z-statistic | p.value |
| --- | --- | --- | --- | --- |
| (Intercept) | -5.748 | 0.632 | -9.095 | 9.488x10^-20^ |
| Distance (km) | -0.055 | 0.024 | -2.312 | 0.021 |
| Direction (-1, 0, 1) | -0.089 | 0.216 | -0.412 | 0.681 |
| Destination habitat size | 0.002 | 0.000 | 6.557 | 5.503x10^-11^ |
| Source habitat size | 0.001 | 0.000 | 5.223 | 1.763 x10^-7^ |
| SWM season | 1.711 | 0.491 | 3.484 | 4.947 x10^-4^ |

**Table S7 Custom barcodes**

The 48 custom barcodes used to multiplex pooled samples during ddRADSeq.

| Name | Oligonucleotide sequence |
| --- | --- |
| P1.1 - SbfI-AAACAC | ACACTCTTTCCCTACACGACGCTCTTCCGATCTAAACACTGCA |
| P1.2 - SbfI-AAACAC | /5phos/GTGTTTAGATCGGAAGAGCGTCGTGTAGGGAAAGAGTGT |
| P1.1 - SbfI-AAACGA | ACACTCTTTCCCTACACGACGCTCTTCCGATCTAAACGATGCA |
| P1.2 - SbfI-AAACGA | /5phos/TCGTTTAGATCGGAAGAGCGTCGTGTAGGGAAAGAGTGT |
| P1.1 - SbfI-AAAGTC | ACACTCTTTCCCTACACGACGCTCTTCCGATCTAAAGTCTGCA |
| P1.2 - SbfI-AAAGTC | /5phos/GACTTTAGATCGGAAGAGCGTCGTGTAGGGAAAGAGTGT |
| P1.1 - SbfI-AACGGT | ACACTCTTTCCCTACACGACGCTCTTCCGATCTAACGGTTGCA |
| P1.2 - SbfI-AACGGT | /5phos/ACCGTTAGATCGGAAGAGCGTCGTGTAGGGAAAGAGTGT |
| P1.1 - SbfI-AACTTC | ACACTCTTTCCCTACACGACGCTCTTCCGATCTAACTTCTGCA |
| P1.2 - SbfI-AACTTC | /5phos/GAAGTTAGATCGGAAGAGCGTCGTGTAGGGAAAGAGTGT |
| P1.1 - SbfI-TGCTCA | ACACTCTTTCCCTACACGACGCTCTTCCGATCTTGCTCATGCA |
| P1.2 - SbfI-TGCTCA | /5phos/TGAGCAAGATCGGAAGAGCGTCGTGTAGGGAAAGAGTGT |
| P1.2 - SbfI-TGCTCA | ACACTCTTTCCCTACACGACGCTCTTCCGATCTAAGAACTGCA |
| P1.1 - SbfI-AAGAAC | /5phos/GTTCTTAGATCGGAAGAGCGTCGTGTAGGGAAAGAGTGT |
| P1.2 - SbfI-AAGAAC | ACACTCTTTCCCTACACGACGCTCTTCCGATCTAATGTGTGCA |
| P1.1 - SbfI-AATGTG | /5phos/CACATTAGATCGGAAGAGCGTCGTGTAGGGAAAGAGTGT |
| P1.2 - SbfI-AATGTG | ACACTCTTTCCCTACACGACGCTCTTCCGATCTACATGTTGCA |
| P1.1 - SbfI-ACATGT | /5phos/ACATGTAGATCGGAAGAGCGTCGTGTAGGGAAAGAGTGT |
| P1.2 - SbfI-ACATGT | ACACTCTTTCCCTACACGACGCTCTTCCGATCTACCAAATGCA |
| P1.1 - SbfI-ACCAAA | /5phos/TTTGGTAGATCGGAAGAGCGTCGTGTAGGGAAAGAGTGT |
| P1.2 - SbfI-ACCAAA | ACACTCTTTCCCTACACGACGCTCTTCCGATCTACGATATGCA |
| P1.1 - SbfI-ACGATA | /5phos/TATCGTAGATCGGAAGAGCGTCGTGTAGGGAAAGAGTGT |
| P1.2 - SbfI-ACGATA | ACACTCTTTCCCTACACGACGCTCTTCCGATCTACGTTTTGCA |
| P1.1 - SbfI-ACGTTT | /5phos/AAACGTAGATCGGAAGAGCGTCGTGTAGGGAAAGAGTGT |
| P1.2 - SbfI-ACGTTT | ACACTCTTTCCCTACACGACGCTCTTCCGATCTACTAGGTGCA |
| P1.1 - SbfI-ACTAGG | /5phos/CCTAGTAGATCGGAAGAGCGTCGTGTAGGGAAAGAGTGT |
| P1.2 - SbfI-ACTAGG | ACACTCTTTCCCTACACGACGCTCTTCCGATCTACTCCATGCA |
| P1.1 - SbfI-ACTCCA | /5phos/TGGAGTAGATCGGAAGAGCGTCGTGTAGGGAAAGAGTGT |
| P1.2 - SbfI-ACTCCA | ACACTCTTTCCCTACACGACGCTCTTCCGATCTAGACTCTGCA |
| P1.1 - SbfI-AGACTC | /5phos/GAGTCTAGATCGGAAGAGCGTCGTGTAGGGAAAGAGTGT |
| P1.2 - SbfI-AGACTC | ACACTCTTTCCCTACACGACGCTCTTCCGATCTAGCATTTGCA |
| P1.1 - SbfI-AGCATT | /5phos/AATGCTAGATCGGAAGAGCGTCGTGTAGGGAAAGAGTGT |
| P1.2 - SbfI-AGCATT | ACACTCTTTCCCTACACGACGCTCTTCCGATCTAGGAGATGCA |
| P1.1 - SbfI-AGGAGA | /5phos/TCTCCTAGATCGGAAGAGCGTCGTGTAGGGAAAGAGTGT |
| P1.2 - SbfI-AGGAGA | ACACTCTTTCCCTACACGACGCTCTTCCGATCTAGTAAGTGCA |
| P1.1 - SbfI-AGTAAG | /5phos/CTTACTAGATCGGAAGAGCGTCGTGTAGGGAAAGAGTGT |
| P1.2 - SbfI-AGTAAG | ACACTCTTTCCCTACACGACGCTCTTCCGATCTAGTCCTTGCA |
| P1.1 - SbfI-AGTCCT | /5phos/AGGACTAGATCGGAAGAGCGTCGTGTAGGGAAAGAGTGT |
| P1.2 - SbfI-AGTCCT | ACACTCTTTCCCTACACGACGCTCTTCCGATCTAGTTACTGCA |
| P1.1 - SbfI-AGTTAC | /5phos/GTAACTAGATCGGAAGAGCGTCGTGTAGGGAAAGAGTGT |
| P1.2 - SbfI-AGTTAC | ACACTCTTTCCCTACACGACGCTCTTCCGATCTATAACCTGCA |
| P1.1 - SbfI-ATAACC | /5phos/GGTTATAGATCGGAAGAGCGTCGTGTAGGGAAAGAGTGT |
| P1.2 - SbfI-ATAACC | ACACTCTTTCCCTACACGACGCTCTTCCGATCTATCGCATGCA |
| P1.1 - SbfI-ATCGCA | /5phos/TGCGATAGATCGGAAGAGCGTCGTGTAGGGAAAGAGTGT |
| P1.2 - SbfI-ATCGCA | ACACTCTTTCCCTACACGACGCTCTTCCGATCTATCTCGTGCA |
| P1.1 - SbfI-ATCTCG | /5phos/CGAGATAGATCGGAAGAGCGTCGTGTAGGGAAAGAGTGT |
| P1.2 - SbfI-ATCTCG | ACACTCTTTCCCTACACGACGCTCTTCCGATCTATGGAGTGCA |
| P1.1 - SbfI-ATGGAG | /5phos/CTCCATAGATCGGAAGAGCGTCGTGTAGGGAAAGAGTGT |
| P1.2 - SbfI-ATGGAG | ACACTCTTTCCCTACACGACGCTCTTCCGATCTATGTCCTGCA |
| P1.1 - SbfI-ATGTCC | /5phos/GGACATAGATCGGAAGAGCGTCGTGTAGGGAAAGAGTGT |
| P1.2 - SbfI-ATGTCC | ACACTCTTTCCCTACACGACGCTCTTCCGATCTCAATTGTGCA |
| P1.1 - SbfI-CAATTG | /5phos/CAATTGAGATCGGAAGAGCGTCGTGTAGGGAAAGAGTGT |
| P1.2 - SbfI-CAATTG | ACACTCTTTCCCTACACGACGCTCTTCCGATCTCAGAGTTGCA |
| P1.1 - SbfI-CAGAGT | /5phos/ACTCTGAGATCGGAAGAGCGTCGTGTAGGGAAAGAGTGT |
| P1.2 - SbfI-CAGAGT | ACACTCTTTCCCTACACGACGCTCTTCCGATCTCATCAGTGCA |
| P1.1 - SbfI-CATCAG | /5phos/CTGATGAGATCGGAAGAGCGTCGTGTAGGGAAAGAGTGT |
| P1.2 - SbfI-CATCAG | ACACTCTTTCCCTACACGACGCTCTTCCGATCTCATCTCTGCA |
| P1.1 - SbfI-CATCTC | /5phos/GAGATGAGATCGGAAGAGCGTCGTGTAGGGAAAGAGTGT |
| P1.2 - SbfI-CATCTC | ACACTCTTTCCCTACACGACGCTCTTCCGATCTCCACTTTGCA |
| P1.1 - SbfI-CCACTT | /5phos/AAGTGGAGATCGGAAGAGCGTCGTGTAGGGAAAGAGTGT |
| P1.2 - SbfI-CCACTT | ACACTCTTTCCCTACACGACGCTCTTCCGATCTCCCATATGCA |
| P1.1 - SbfI-CCCATA | /5phos/TATGGGAGATCGGAAGAGCGTCGTGTAGGGAAAGAGTGT |
| P1.2 - SbfI-CCCATA | ACACTCTTTCCCTACACGACGCTCTTCCGATCTCCTGAATGCA |
| P1.1 - SbfI-CCTGAA | /5phos/TTCAGGAGATCGGAAGAGCGTCGTGTAGGGAAAGAGTGT |
| P1.2 - SbfI-CCTGAA | ACACTCTTTCCCTACACGACGCTCTTCCGATCTCGAAACTGCA |
| P1.1 - SbfI-CGAAAC | /5phos/GTTTCGAGATCGGAAGAGCGTCGTGTAGGGAAAGAGTGT |
| P1.2 - SbfI-CGAAAC | ACACTCTTTCCCTACACGACGCTCTTCCGATCTCGAATGTGCA |
| P1.1 - SbfI-CGAATG | /5phos/CATTCGAGATCGGAAGAGCGTCGTGTAGGGAAAGAGTGT |
| P1.2 - SbfI-CGAATG | ACACTCTTTCCCTACACGACGCTCTTCCGATCTGACGTTTGCA |
| P1.1 - SbfI-GACGTT | /5phos/AACGTCAGATCGGAAGAGCGTCGTGTAGGGAAAGAGTGT |
| P1.2 - SbfI-GACGTT | ACACTCTTTCCCTACACGACGCTCTTCCGATCTGATACATGCA |
| P1.1 - SbfI-GATACA | /5phos/TGTATCAGATCGGAAGAGCGTCGTGTAGGGAAAGAGTGT |
| P1.2 - SbfI-GATACA | ACACTCTTTCCCTACACGACGCTCTTCCGATCTGCAGAATGCA |
| P1.1 - SbfI-GCAGAA | /5phos/TTCTGCAGATCGGAAGAGCGTCGTGTAGGGAAAGAGTGT |
| P1.2 - SbfI-GCAGAA | ACACTCTTTCCCTACACGACGCTCTTCCGATCTGGGATATGCA |
| P1.1 - SbfI-GGGATA | /5phos/TATCCCAGATCGGAAGAGCGTCGTGTAGGGAAAGAGTGT |
| P1.2 - SbfI-GGGATA | ACACTCTTTCCCTACACGACGCTCTTCCGATCTGGTGAATGCA |
| P1.1 - SbfI-GGTGAA | /5phos/TTCACCAGATCGGAAGAGCGTCGTGTAGGGAAAGAGTGT |
| P1.2 - SbfI-GGTGAA | ACACTCTTTCCCTACACGACGCTCTTCCGATCTGTAGCTTGCA |
| P1.1 - SbfI-GTAGCT | /5phos/AGCTACAGATCGGAAGAGCGTCGTGTAGGGAAAGAGTGT |
| P1.2 - SbfI-GTAGCT | ACACTCTTTCCCTACACGACGCTCTTCCGATCTGTCTATTGCA |
| P1.1 - SbfI-GTCTAT | /5phos/ATAGACAGATCGGAAGAGCGTCGTGTAGGGAAAGAGTGT |
| P1.2 - SbfI-GTCTAT | ACACTCTTTCCCTACACGACGCTCTTCCGATCTGTTCAGTGCA |
| P1.1 - SbfI-GTTCAG | /5phos/CTGAACAGATCGGAAGAGCGTCGTGTAGGGAAAGAGTGT |
| P1.2 - SbfI-GTTCAG | ACACTCTTTCCCTACACGACGCTCTTCCGATCTTAAGACTGCA |
| P1.1 - SbfI-TAAGAC | /5phos/GTCTTAAGATCGGAAGAGCGTCGTGTAGGGAAAGAGTGT |
| P1.2 - SbfI-TAAGAC | ACACTCTTTCCCTACACGACGCTCTTCCGATCTTACCAGTGCA |
| P1.1 - SbfI-TACCAG | /5phos/CTGGTAAGATCGGAAGAGCGTCGTGTAGGGAAAGAGTGT |
| P1.2 - SbfI-TACCAG | ACACTCTTTCCCTACACGACGCTCTTCCGATCTTCAATCTGCA |
| P1.1 - SbfI-TCAATC | /5phos/GATTGAAGATCGGAAGAGCGTCGTGTAGGGAAAGAGTGT |
| P1.2 - SbfI-TCAATC | ACACTCTTTCCCTACACGACGCTCTTCCGATCTTCCAAATGCA |
| P1.1 - SbfI-TCCAAA | /5phos/TTTGGAAGATCGGAAGAGCGTCGTGTAGGGAAAGAGTGT |
| P1.2 - SbfI-TCCAAA | ACACTCTTTCCCTACACGACGCTCTTCCGATCTTCTGCTTGCA |
| P1.1 - SbfI-TCTGCT | /5phos/AGCAGAAGATCGGAAGAGCGTCGTGTAGGGAAAGAGTGT |
| P1.2 - SbfI-TCTGCT | ACACTCTTTCCCTACACGACGCTCTTCCGATCTTCTTAGTGCA |
| P1.1 - SbfI-TCTTAG | /5phos/CTAAGAAGATCGGAAGAGCGTCGTGTAGGGAAAGAGTGT |

**Table S8 PCR Primers**

Multiplexing PCR primers used to amplify DNA during ddRADseq.

| Primer | Sequence (5’ to 3’) |
| --- | --- |
| PCR1 | AATGATACGGCGACCACCGAGATCTACACTCTTTCCCTACACGACG |
| PCR2 Index 2 | CAAGCAGAAGACGGCATACGAGATCGTGATGTGACTGGAGTTCAGACGTGTGC |
| PCR2 Index 3 | CAAGCAGAAGACGGCATACGAGATGCCTAAGTGACTGGAGTTCAGACGTGTGC |
| PCR2 Index 4 | CAAGCAGAAGACGGCATACGAGATTGGTCAGTGACTGGAGTTCAGACGTGTGC |
| PCR2 Index 5 | CAAGCAGAAGACGGCATACGAGATCACTGTGTGACTGGAGTTCAGACGTGTGC |
| PCR2 Index 6 | CAAGCAGAAGACGGCATACGAGATATTGGCGTGACTGGAGTTCAGACGTGTGC |
| PCR2 Index 7 | CAAGCAGAAGACGGCATACGAGATGATCTGGTGACTGGAGTTCAGACGTGTGC |
| PCR2 Index 8 | CAAGCAGAAGACGGCATACGAGATTCAAGTGTGACTGGAGTTCAGACGTGTGC |
| PCR2 Index 9 | CAAGCAGAAGACGGCATACGAGATCTGATCGTGACTGGAGTTCAGACGTGTGC |
| PCR2 Index 10 | CAAGCAGAAGACGGCATACGAGATAAGCTAGTGACTGGAGTTCAGACGTGTGC |
| PCR2 Index 11 | CAAGCAGAAGACGGCATACGAGATGTAGCCGTGACTGGAGTTCAGACGTGTGC |
| PCR2 Index 12 | CAAGCAGAAGACGGCATACGAGATTACAAGGTGACTGGAGTTCAGACGTGTGC |

**Table S9** **Sequencing Summary**

Summary of all sequencing runs on an Illumina 2500 at the Princeton Core Facility (Princeton, NJ, USA).

| Sequencing run | Number of samples | Total reads | Reads passing quality filters |
| --- | --- | --- | --- |
| SEQ03 | 192 | 248189468 | 213307370 |
| SEQ04 | 192 | 251852650 | 180383111 |
| SEQ05 | 192 | 251297377 | 182283859 |
| SEQ07 | 192 | 175059438 | 158606770 |
| SEQ08 | 192 | 199919992 | 177884814 |
| SEQ09 | 192 | 171814118 | 141309748 |
| SEQ12 | 192 | 150572315 | 134418274 |
| SEQ13 | 192 | 216158456 | 207836331 |
| SEQ15 | 192 | 237201902 | 227519254 |
| SEQ16 | 192 | 214609013 | 200476806 |
| SEQ17 | 192 | 198878655 | 191839340 |
| SEQ28 | 576 | 207926899 | 203002503 |
| SEQ29 | 576 | 225203742 | 211426551 |
| SEQ30 | 576 | 228316545 | 217444014 |
| SEQ31 | 576 | 125190259 | 117312441 |

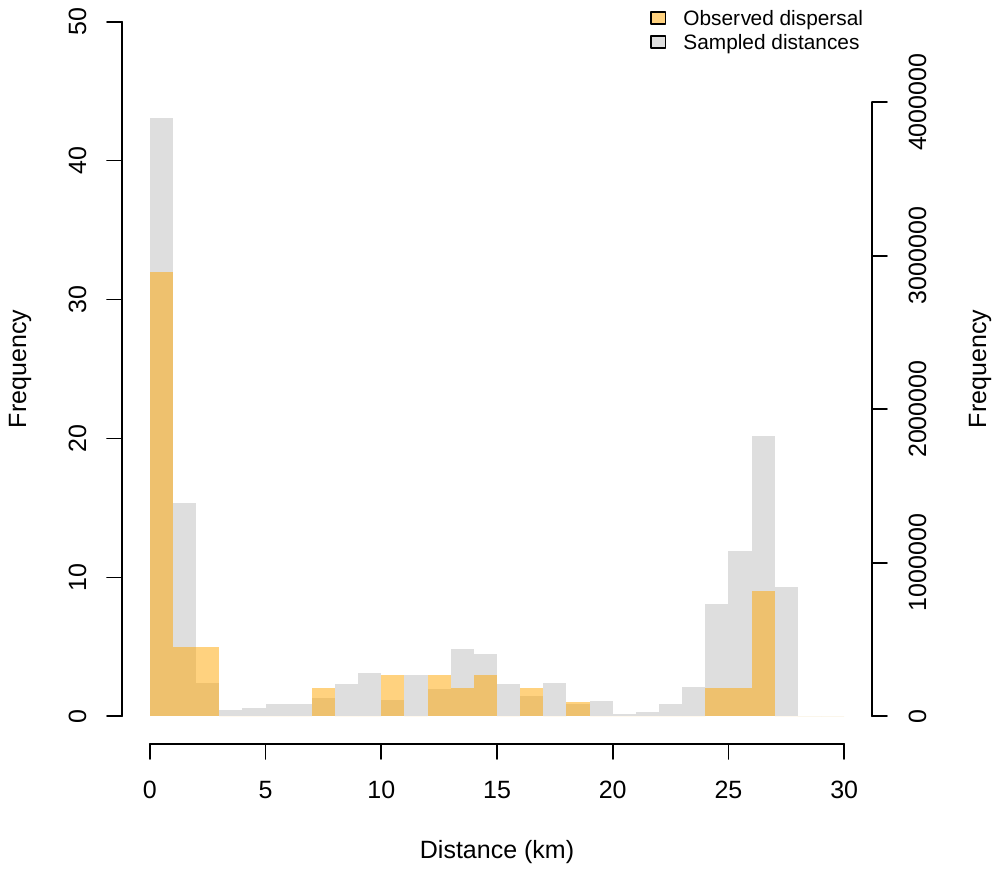

**Figure S1** Frequency of all observed dispersal events at a given distance (orange bars, left hand axis) compared to the frequency of all sampling distances (gray bars, right hand axis). The shape of the sampling distance distribution is determined by the geographic layout of sites and the sampling effort at sites.

**Figure S2** Bivariate 95% CI likelihood surfaces for the annual kernel fits of *k* (scale parameter) and θ (shape parameter). Colors represent the different years.

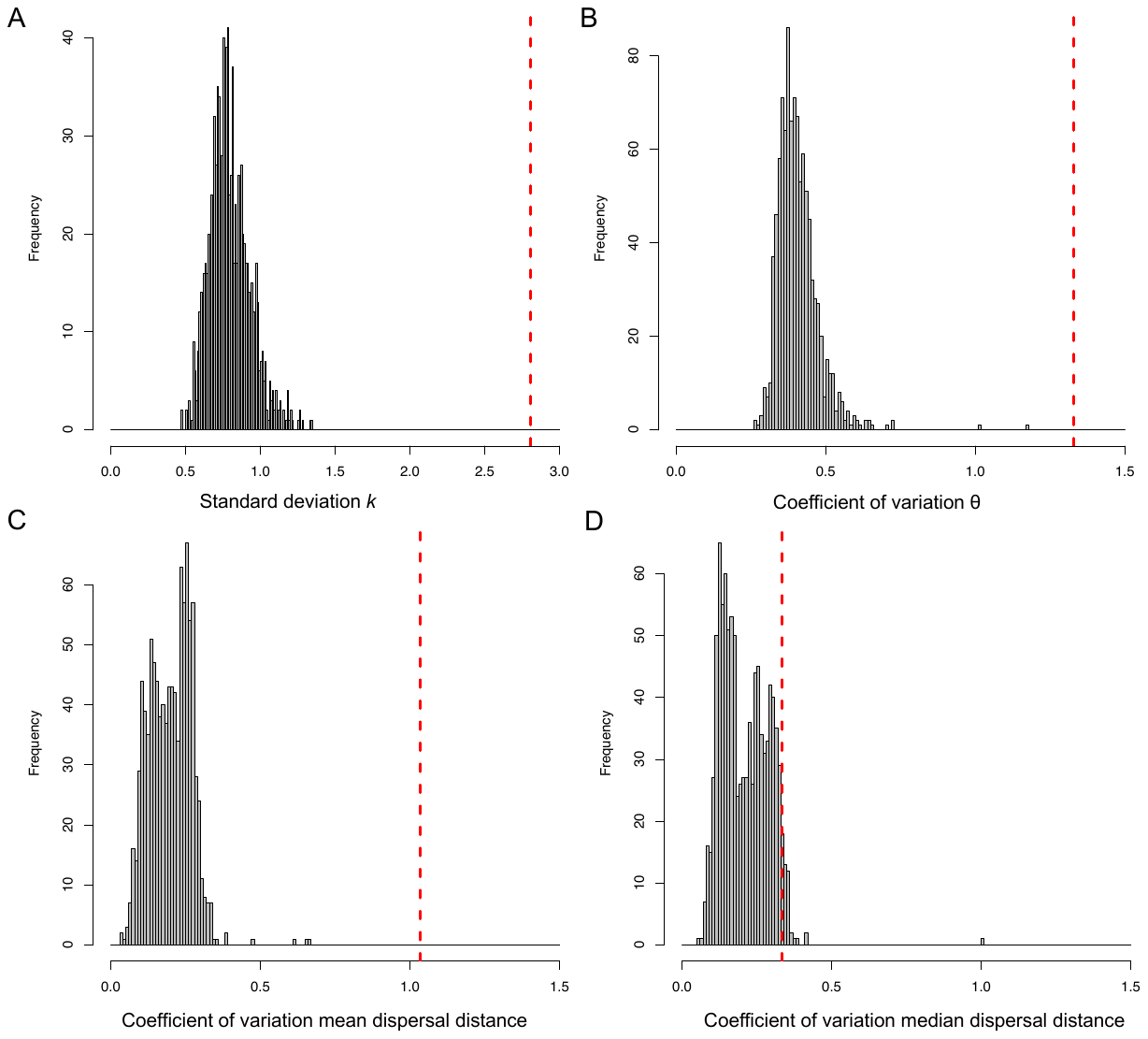

**Figure S3** A) The empirical estimate (red line) of the interannual standard deviation of *k* (2.80) is greater than 0.999 of the null distribution (gray bars). B) Empirical estimate (red line) of the interannual coefficient of variation of θ (CV=1.33) is greater than 0.999 of the null distribution (gray bars). C) Empirical estimate (red line) of the interannual CV of mean dispersal distance (1.03) is greater than 0.999 of the null distribution (gray bars). D) Empirical estimate (red line) of the interannual CV of median dispersal distance (0.34) is greater than 0.959 of the null distribution (gray bars).

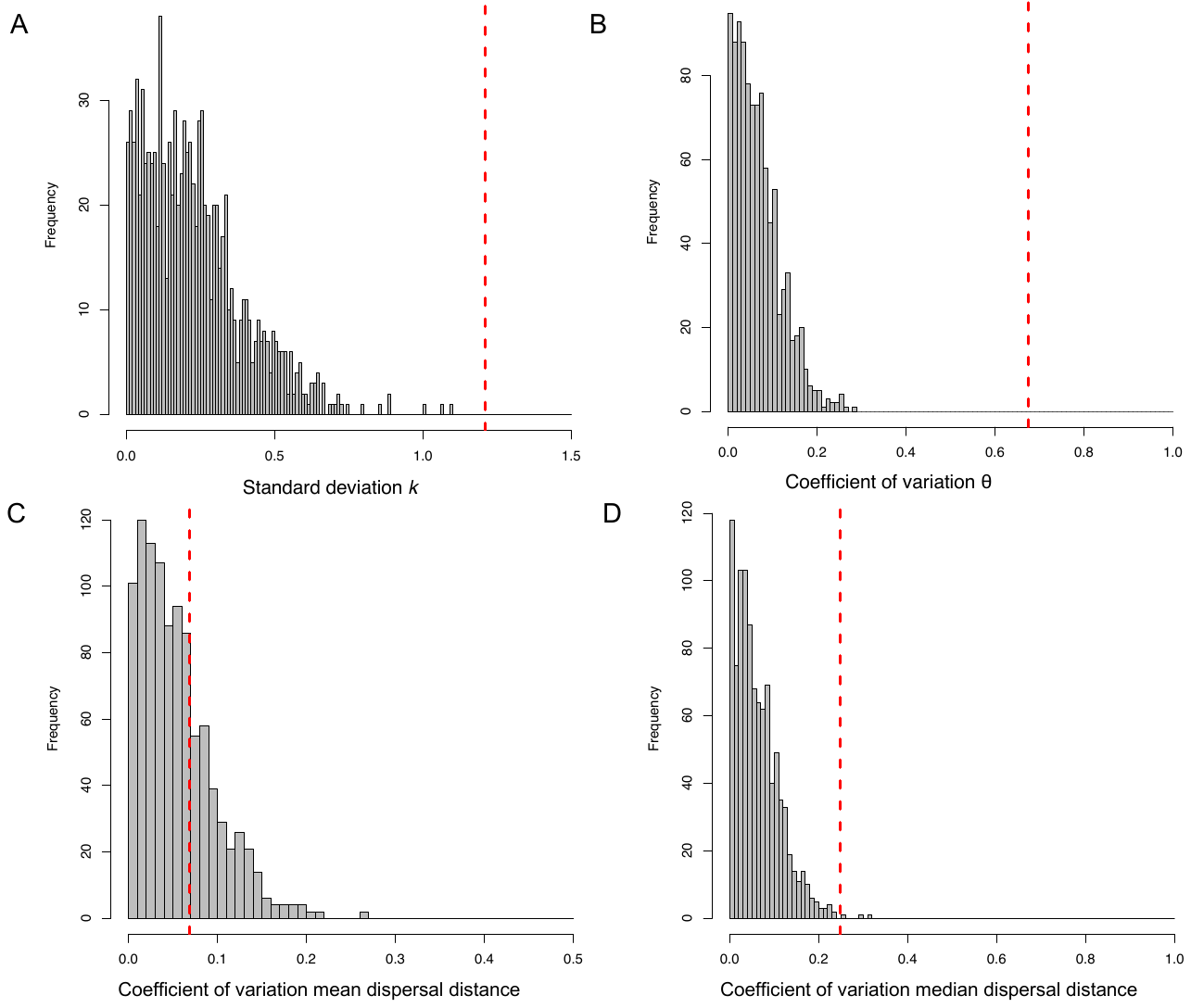

**Figure S4** A) The empirical estimate (red line) of the seasonal standard deviation of *k* (1.21) is greater than 0.999 of the null distribution (gray bars). B) Empirical estimate (red line) of the seasonal coefficient of variation of θ (CV=0.67) is greater than 0.999 of the null distribution (gray bars). C) Empirical estimate (red line) of the seasonal CV of mean dispersal distance (0.07) is greater than 0.681 of the null distribution (gray bars). D) Empirical estimate (red line) of the seasonal CV of median dispersal distance (0.25) is greater than 0.997 of the null distribution (gray bars).

­­­
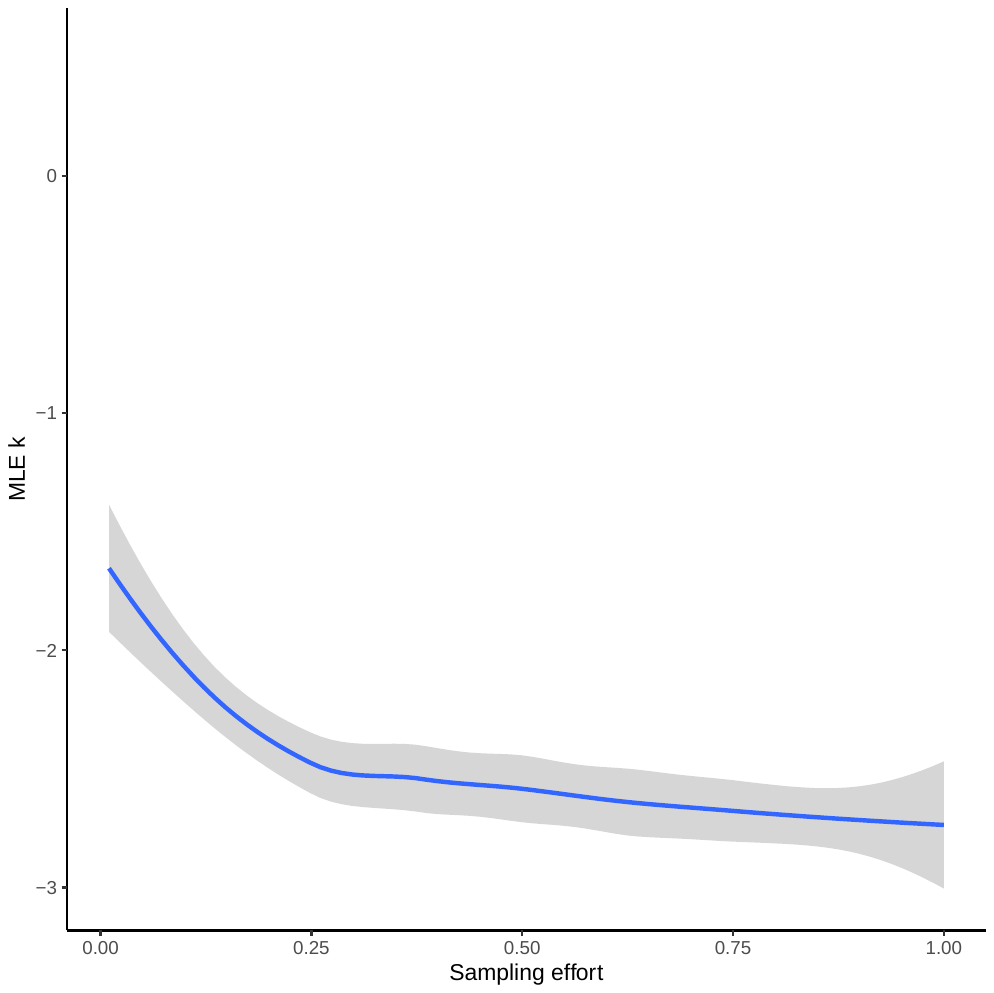

**Figure S5** Results of simulations exploring sensitivity of kernel parameter *k* to the proportion of individuals sampled (from 0.01 to 1.0, *n* = 100) using the data across all years.

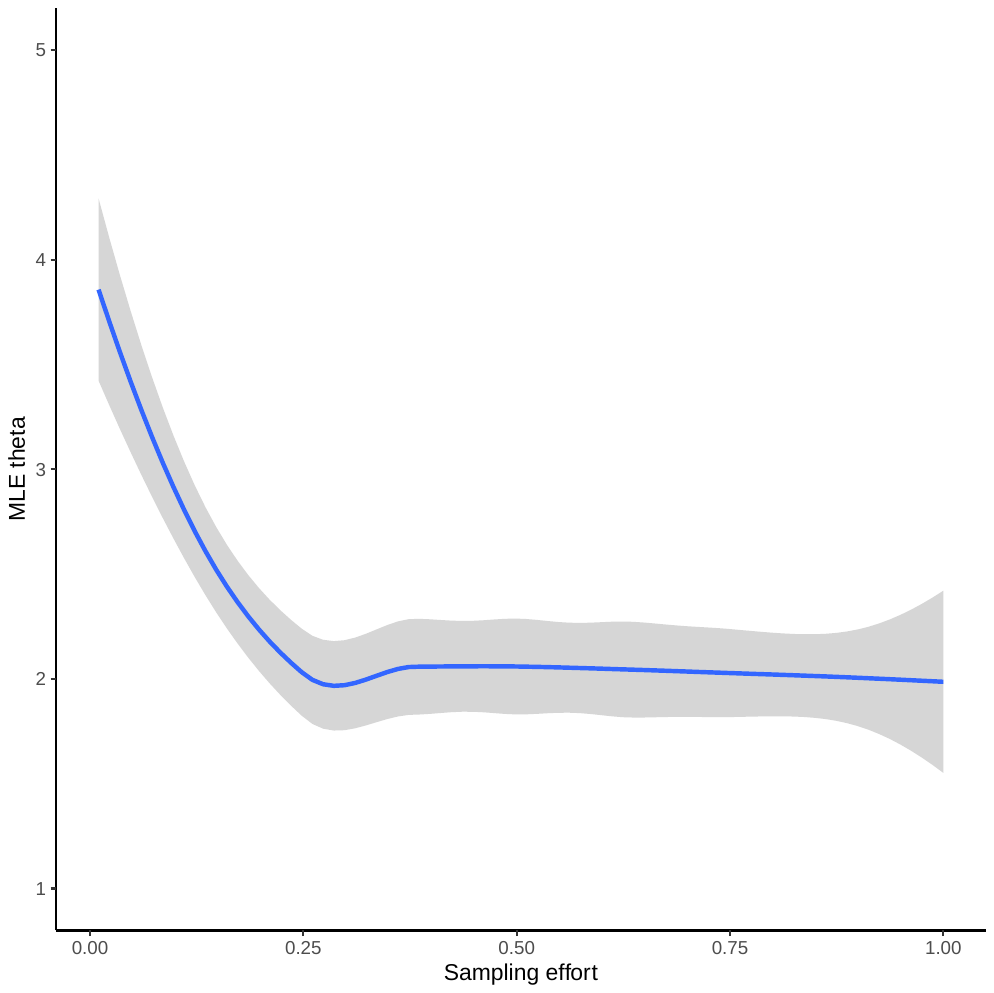

**Figure S6** Results of simulations exploring sensitivity of kernel parameter θ to sampling effort, as in the proportion of individuals sampled (from 0.01 to 1.0, *n* = 100) using the data across all years.

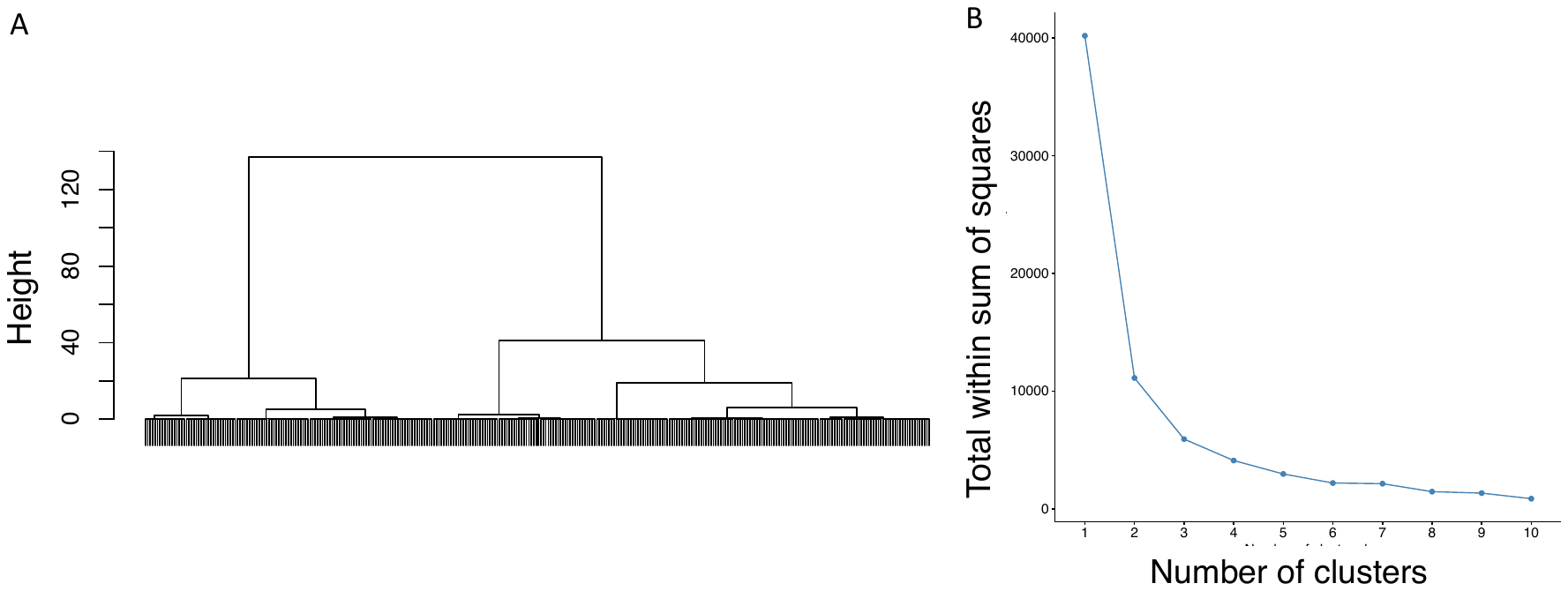

**Figure S7** A) Cluster dendrogram illustrating the three groups identified by Ward’s hierarchical clustering method applied to the dispersal route correlation matrix. Each terminal branch represents a dispersal route. B) The relationship between total within-cluster sum of squares and number of clusters. The addition of clusters beyond three showed relatively little improvement in minimizing within-cluster sum of squares.
